# A membrane-depolarizing toxin substrate of the *Staphylococcus aureus* Type VII secretion system mediates intra-species competition

**DOI:** 10.1101/443630

**Authors:** Fatima R. Ulhuq, Margarida C. Gomes, Gina Duggan, Manman Guo, Chriselle Mendonca, Grant Buchanan, James D. Chalmers, Zhenping Cao, Holger Kneuper, Sarah Murdoch, Sarah Thomson, Henrik Strahl, Matthias Trost, Serge Mostowy, Tracy Palmer

**Affiliations:** Molecular and Cellular Microbiology Theme, Newcastle University Biosciences Institute, Newcastle University, Newcastle upon Tyne, NE2 4HH, UK; Section of Microbiology, MRC Centre for Molecular Bacteriology and Infection, Imperial College London, Armstrong Road, London SW7 2AZ, UK; Department of Infection Biology, London School of Hygiene & Tropical Medicine, Keppel Street, London WC1E 7HT, UK; MRC Protein Phosphorylation and Ubiquitylation Unit, School of Life Sciences, University of Dundee, Dundee DD1 5EH, UK; Division of Molecular Microbiology, School of Life Sciences, University of Dundee, Dundee DD1 5EH, UK; Division of Molecular and Clinical Medicine, University of Dundee, Dundee, DD1 9SY, UK; Biological Services, School of Life Sciences, University of Dundee, Dundee DD1 5EH, UK; Newcastle University Biosciences Institute, Newcastle University, Newcastle upon Tyne, NE2 4HH, UK

**Keywords:** Type VII secretion system, *Staphylococcus aureus*, zebrafish, membrane-depolarizing toxin, bacterial competition

## Abstract

The type VII protein secretion system (T7SS) is conserved across *Staphylococcus aureus* strains and plays important roles in virulence and interbacterial competition. To date only one T7SS substrate protein, encoded in a subset of *S. aureus* genomes, has been functionally characterized. Here, using an unbiased proteomic approach, we identify TspA as a further T7SS substrate. TspA is encoded distantly from the T7SS gene cluster and is found across all *S. aureus* strains as well as in *Listeria* and Enterococci. Heterologous expression of TspA from *S. aureus* strain RN6390 indicates its C-terminal domain is toxic when targeted to the *Escherichia coli* periplasm and that it depolarizes the cytoplasmic membrane. The membrane depolarizing activity is alleviated by co-production of the membrane-bound TsaI immunity protein, which is encoded adjacent to *tspA* on the *S. aureus* chromosome. Using a zebrafish hindbrain ventricle infection model, we demonstrate that the T7SS of strain RN6390 promotes bacterial replication *in vivo*, and deletion of *tspA* leads to increased bacterial clearance. The toxin domain of TspA is highly polymorphic and *S. aureus* strains encode multiple *tsaI* homologues at the *tspA* locus, suggestive of additional roles in intra-species competition. In agreement, we demonstrate TspA-dependent growth inhibition of RN6390 by strain COL in the zebrafish infection model that is alleviated by the presence of TsaI homologues.

**Significance statement:** *Staphylococcus aureus*, a human commensal organism that asymptomatically colonizes the nares, is capable of causing serious disease following breach of the mucosal barrier. *S. aureus* strains encode a Type VII secretion system (T7SS) that is required for virulence in mouse infection models, and some strains also secrete a nuclease toxin by this route that has antibacterial activity. Here we identify TspA, widely found in Staphylococci and other pathogenic bacteria, as a T7 substrate. We show that TspA has membrane-depolarizing activity and that *S. aureus* uses TspA to inhibit the growth of a bacterial competitor *in vivo*.

## Introduction

The Type VII secretion system (T7SS) has been characterized in bacteria of the actinobacteria and firmicutes phyla. In pathogenic mycobacteria the ESX-1 T7SS secretes numerous proteins that are essential for virulence and immune evasion^1^. The Ess T7SS of *Staphylococcus aureus* is also required for pathogenesis in murine models of infection^2-4^, and a longitudinal study of persistent *S. aureus* infection in the airways of a cystic fibrosis patient showed that the *ess* T7SS genes were highly upregulated during a 13 year timespan^5^. It is becoming increasingly apparent, however, that in addition to having anti-eukaryotic activity, the T7SS of firmicutes mediates interbacterial competition^6-8^. Some strains of *S. aureus* secrete a DNA endonuclease toxin, EsaD^6, 9^, that when overproduced leads to growth inhibition of a sensitive *S. aureus* strain^6^. Moreover, *Streptococcus intermedius* exports at least three LXG domain-containing toxins, TelA, TelB and TelC that mediate contact-dependent growth inhibition against a range of Gram-positive species^7^.

A large integral membrane ATPase of the FtsK/SpoIIIE family, termed EssC in firmicutes, is a conserved component of all T7SSs and probably energizes protein secretion as well as forming part of the translocation channel^10-15^. EsxA, a small secreted protein of the WXG100 family, is a further conserved T7 component that is dependent on the T7SS for its translocation across the membrane^2, 16^. In firmicutes three further membrane proteins, EsaA, EssA and EssB, function alongside the EssC ATPase to mediate T7 protein secretion^17, 18^. In *S. aureus* the T7 structural components are encoded at the *ess* locus. In commonly-studied strains including Newman, RN6390 and USA300, the T7 substrates EsxB, EsxC, EsxD and EsaD are encoded immediately downstream of *essC* (Fig 1A) and are co-regulated with the genes coding for machinery components^2, 3, 6, 9, 19, 20^. With the exception of EsaD the biological activities of these substrates are unknown, although mutational studies have suggested that EsxB and EsxC contribute to persistent infection in a murine abscess model^2, 19^.

**Figure 1.**
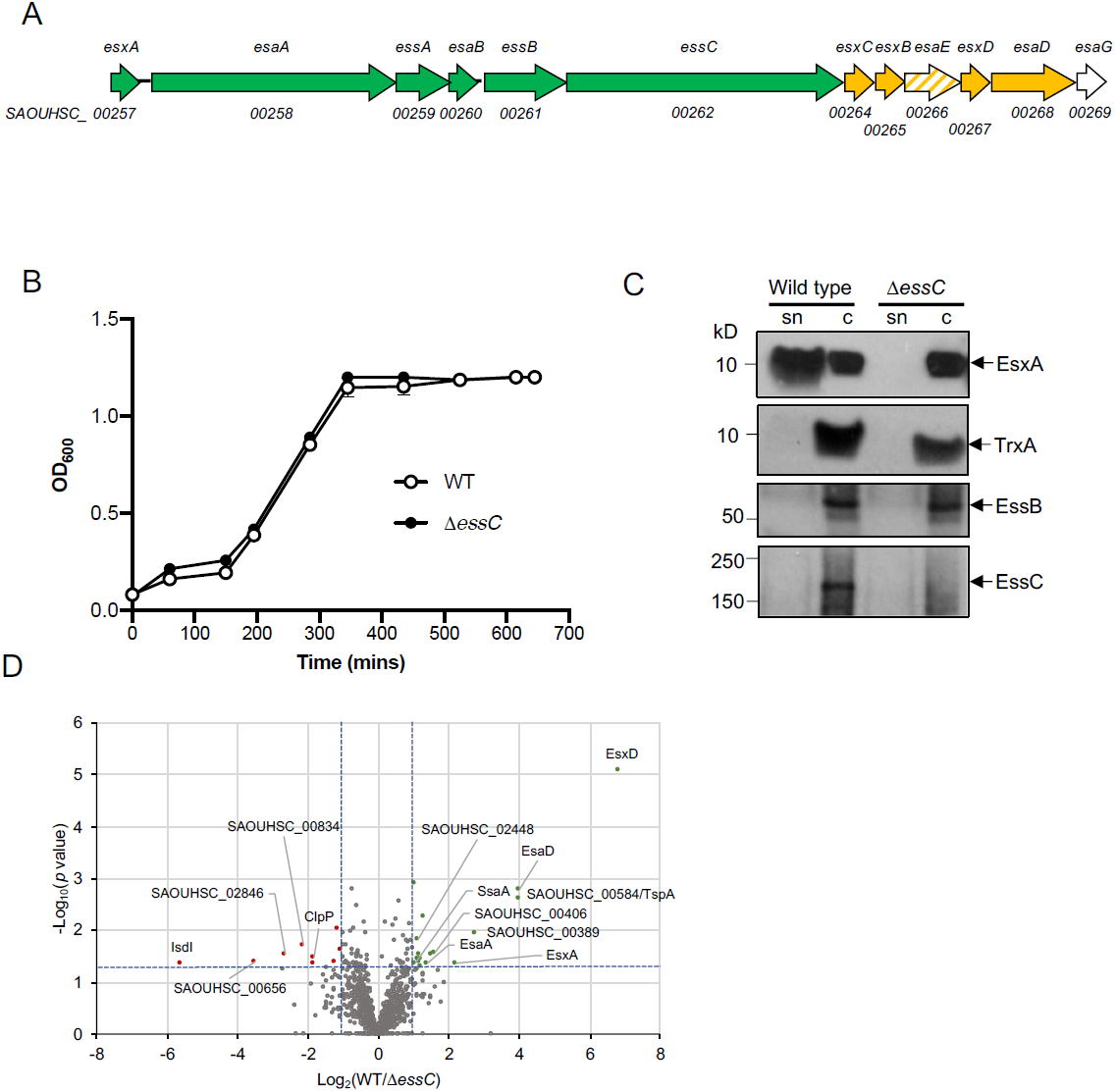
The *S. aureus* RN6390 T7 secretome. A. the *ess* locus in strain NCTC8325 (parent of RN6390). Genes for core components are shaded green, secreted substrates yellow, EsaE (which is co-secreted with EsaD) in hatched shading and the cytoplasmic antitoxin EsaG in white. B. Growth of RN6390 (WT) and the isogenic Δ*essC* strain in RPMI medium. Points show mean +/– SEM (*n* = 3 biological replicates). C. RN6390 (WT) and the Δ*essC* strain cultured in RPMI growth medium to OD_600_ = 1. Samples were separated into supernatant (sn) and cellular (c) fractions (12% bis-Tris gels) and immunoblotted with anti-EsxA, anti-EssB, anti-EssC or anti-TrxA (cytoplasmic protein) antisera. D. Volcano plot of the quantitative proteomic secretome analysis. Each spot represents an individual protein ranked according to its statistical *p*-value (y-axis) and relative abundance ratio (log_2_ fold change). The blue dotted lines represent cut-offs for significance (*p*<0.05; log_2_ fold-change>1).

Despite the *ess* locus forming part of the core *S. aureus* genome, these four substrate proteins are not conserved across *S. aureus* isolates, being found in only approximately 50% of sequenced strains^21^. Furthermore, inactivation of the T7SS in *S. aureus* strain ST398 shows a similar decrease in kidney abscess formation as that seen for T7 mutants in Newman and USA300^2, 4, 22^, despite the fact that recognizable homologues of EsxB, EsxC, EsxD and EsaD are not encoded by this strain^21^. This strongly suggests that there are further *S. aureus* T7 substrates that are yet to be identified. Here we have taken an unbiased approach to identify T7 substrates using quantitative proteomic analysis of culture supernatants from *S. aureus* RN6390 wild type and *essC* strains. We identify a new substrate, TspA that is encoded distantly from the *ess* gene cluster and is found in all sequenced *S. aureus* strains. Further analysis indicates that TspA has a toxic C-terminal domain that depolarizes membranes. Using a zebrafish hindbrain ventricle infection model, we reveal that the T7SS and TspA contribute to both bacterial replication and interbacterial competition *in vivo*.

## Results

### The *S. aureus* RN6390 T7SS secreted proteome

To identify candidate T7 substrates, RN6390 and an isogenic *essC* deletion strain^3^ were cultured in the minimal medium RPMI. Both strains grew identically (Fig 1B) and expressed components of the T7SS, and as expected secretion of EsxA was abolished in the *essC* strain (Fig 1C). Culture supernatants were isolated when cells reached OD_600nm_ of 1, and label-free quantitative proteomics used to assess changes in protein abundance of four biological replicates of each secretome. Following identification of 1170 proteins, 17 proteins showed, with high confidence (p < 0.05, 2-fold change), a decrease in abundance in the secretome of the *essC* strain relative to the RN6390 parent strain (Fig 1D,Table 1; *SI Appendix*, Table S1).

**Table 1.**
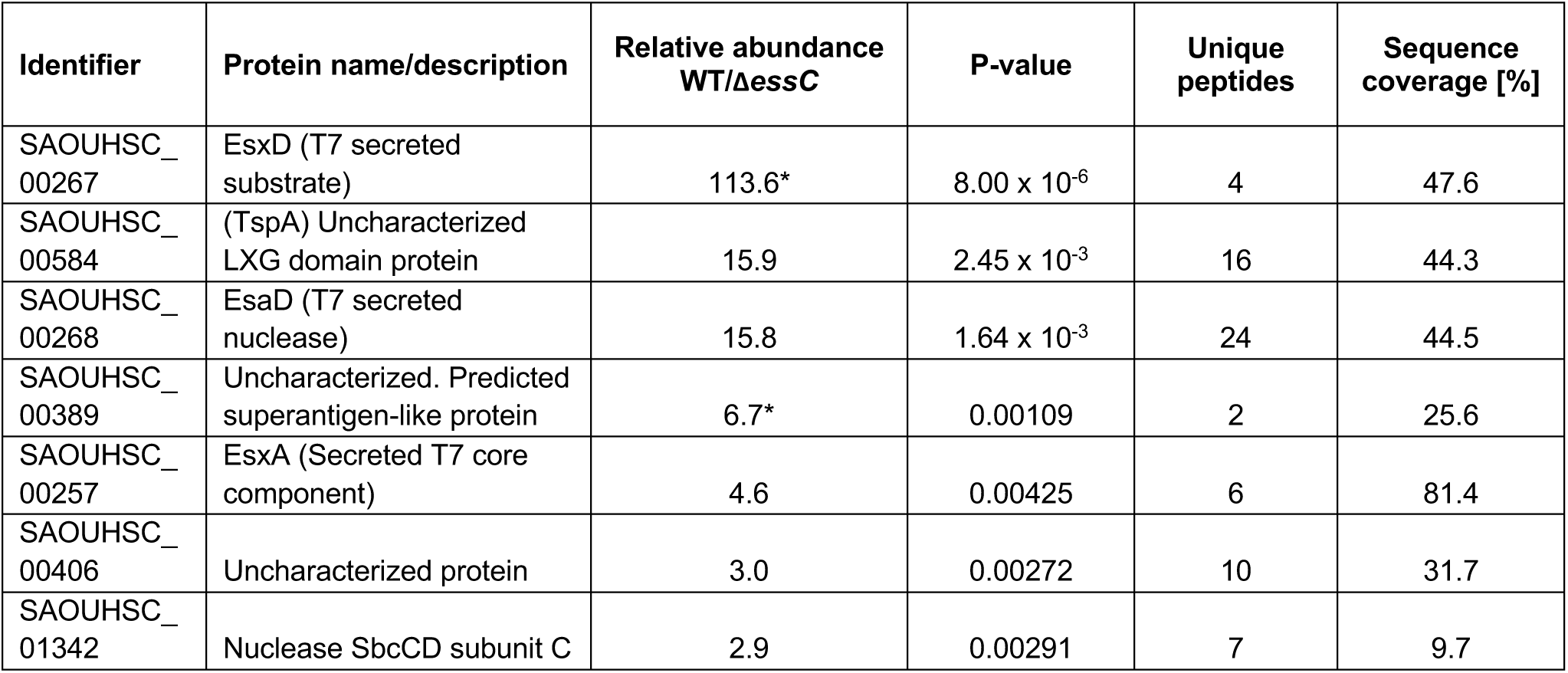
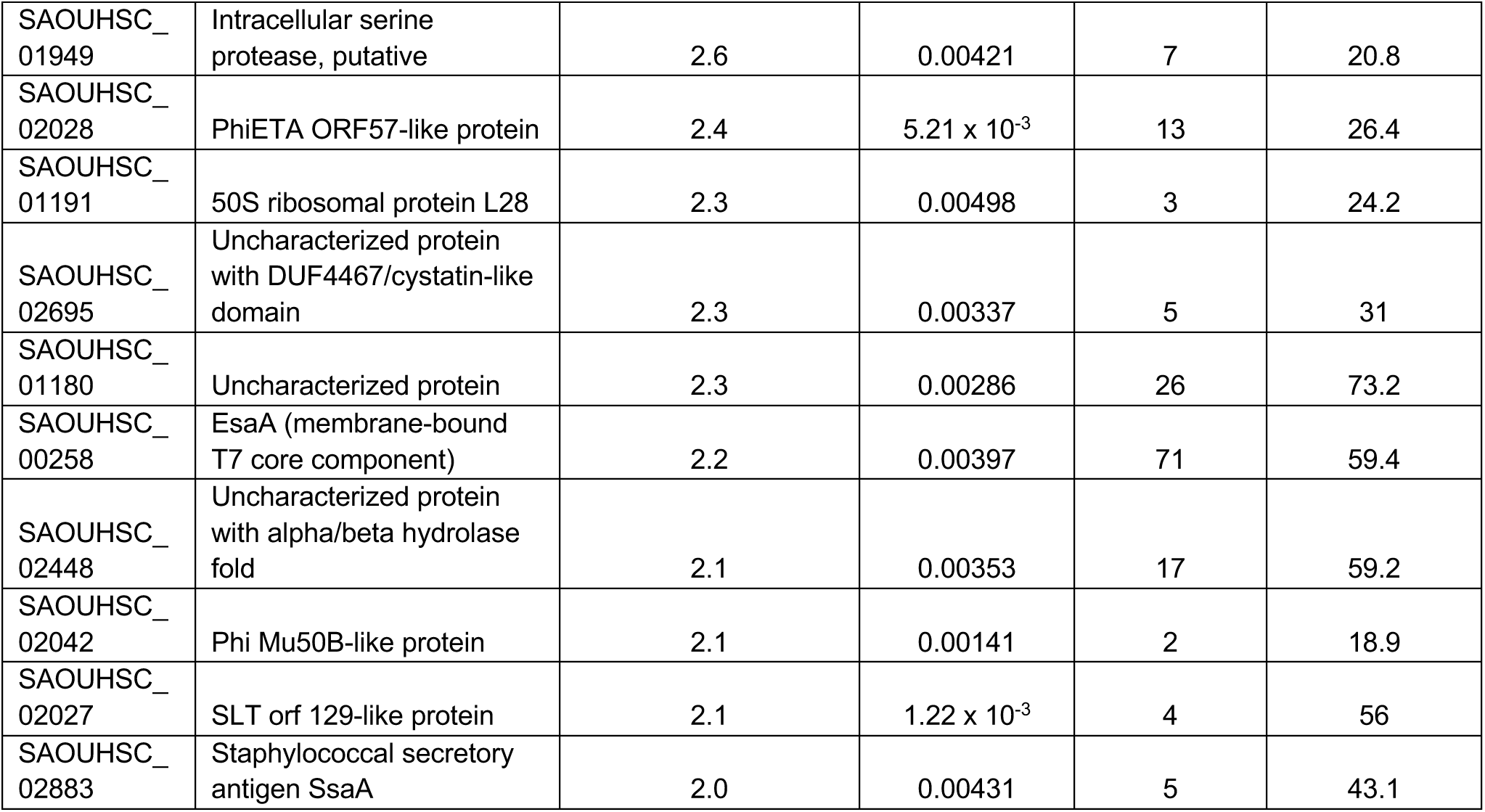
Proteins present in the secretome of RN6390 at an abundance of greater than 2-fold higher than the secretome of the isogenic ΔessC strain. A full list of all of the proteins identified in this analysis is given in *SI Appendix*, Table S1. *not detected in the Δ*essC* secretome.

Proteomic analysis indicated that the secreted core component, EsxA, was significantly reduced in abundance in the *essC* secretome, as expected from the western blot analysis (Fig 1C). Peptides from the membrane-bound T7 component EsaA, which has a large surface-exposed loop^23^, were also less abundant in the supernatant of the *essC* strain, as were EsxD and EsaD, known substrates of the T7SS^6, 9, 20^ (Table 1, *SI Appendix*, Table S1). The EsxC substrate^19^ was also exclusively detected in the supernatant of the wild type strain, but only two EsxC peptides were detected and it did not meet the p < 0.01 cut-off (*SI Appendix*, Table S1). EsxB, another previously identified substrate^2, 24^, and EsaE, which is co-secreted with EsaD^6^ were not detected in any of our analysis.

After EsxD, the protein with the highest relative abundance in the secretome of the wild type strain was the uncharacterized protein SAOUHSC_00584. This protein harbors a predicted LXG domain which is common to some T7SS substrates^7^. Other proteins enriched in the secretome of the wild type strain include a predicted superantigen (SAOUHSC_00389), the secretory antigen SsaA, two predicted membrane-bound lipoproteins (SAOUHSC_01180 and SAOUHSC_02695), two uncharacterized proteins (SAOUHSC_00406 and SAOUHSC_02448) and several predicted cytoplasmic proteins (Table 1). A small number of proteins were found to be enriched in abundance in the *essC* secretome (*SI Appendix*, Table S1), including the heme oxygenase IsdI, which is known to be upregulated when the T7SS is inactivated^25^.

### SAOUHSC_00584/TspA localizes to the membrane dependent on EssC

We next constructed tagged variants of each of SAOUHSC_00389, SAOUHSC_00406, SAOUHSC_00584, SAOUHSC_02448 and SsaA to probe their subcellular locations in the wild type and Δ*essC* strains. C-terminally HA-tagged SAOUHSC_00389, SAOUHSC_02448 and SsaA were secreted into the culture supernatant in an *essC*-independent manner (*SI Appendix*, Fig S1), indicating that these proteins are not substrates of the T7SS and their reduced abundance in the *essC* secretome may arise for pleiotropic reasons. Overproduction of C-terminally Myc-tagged SAOUHSC_00406 caused cell lysis, seen by the presence of TrxA in the supernatant samples (*SI Appendix*, Fig S1B). By contrast, a C-terminally Myc-tagged variant of SAOUHSC_00584 was detected only in the cellular fraction (*SI Appendix*, Fig S1C). To probe the subcellular location of SAOUHSC_00584-Myc, we generated cell wall, membrane and cytoplasmic fractions. Fig 2A shows that the tagged protein localized to the membrane and that it appears to be destabilized by the loss of EssC. SAOUHSC_00584 was subsequently renamed TspA (<u>T</u>ype <u>S</u>even dependent <u>P</u>rotein A).

**Figure 2.**
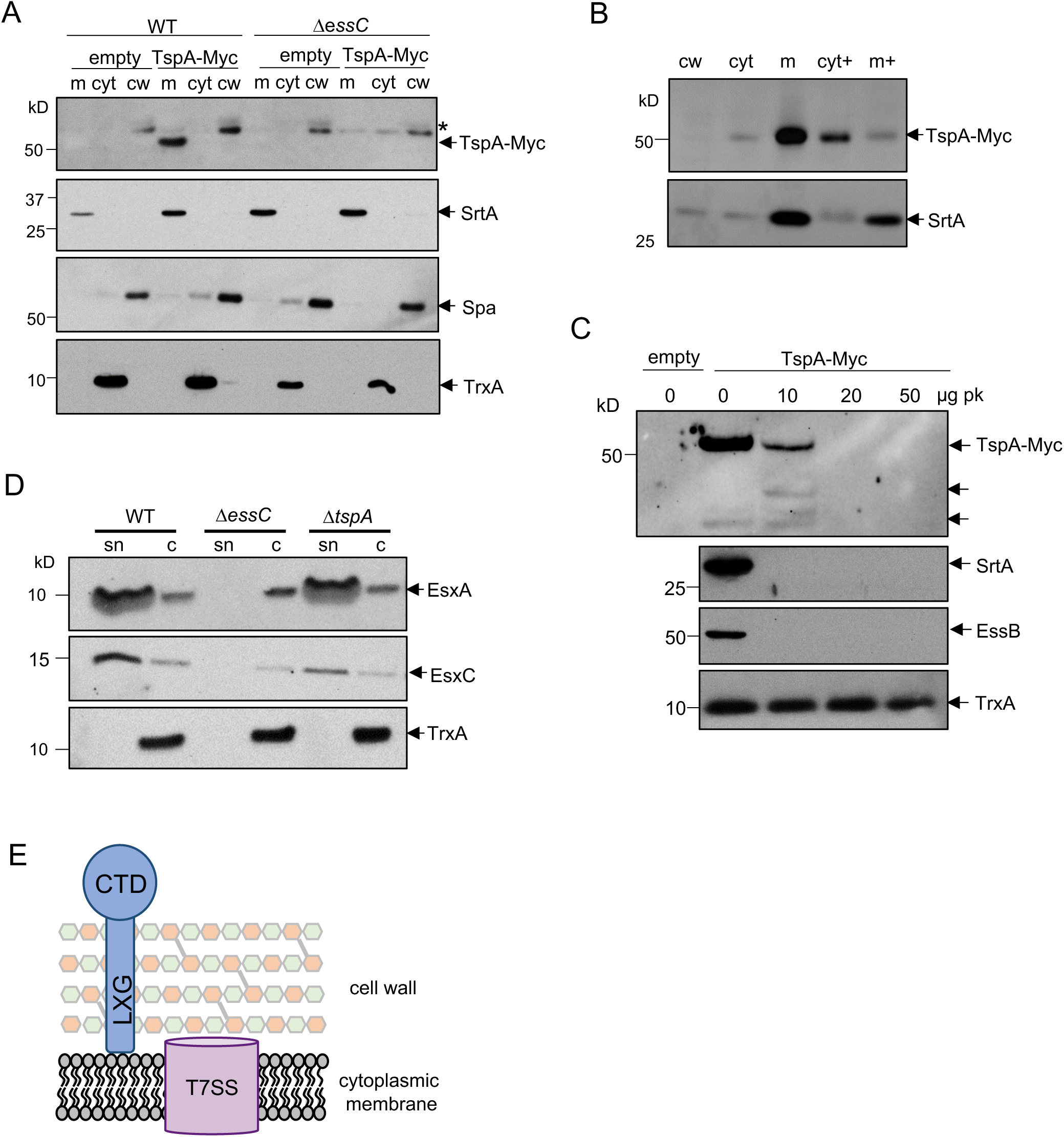
SAOUHSC_00584/TspA is an extracellular peripheral membrane protein. A. RN6390 (WT) and the Δ*essC* strain harbouring pRAB11 (empty) or pRAB11-TspA-Myc were cultured in TSB growth medium. Following induction of plasmid-encoded TspA-Myc production, cells were harvested and fractionated into cell wall (cw), membrane (m) and cytoplasmic (cyt) fractions. Samples were separated (12% bis-Tris gels) and immunoblotted with anti-Myc-HRP anti-TrxA (cytoplasmic protein), anti-Spa (cell wall) or anti-SrtA (membrane) antisera. * ‘non-specific’ cross-reacting band corresponding to Spa. B. Cell extracts from the RN6390 samples in (A) were incubated with 4M urea, membranes were isolated and the urea-treated cytoplasm (cyt+) and membranes (m+) were separated alongside the cell wall and untreated cytoplasm and membrane fractions on a 12% bis-tris gel and immunoblotted with anti-Myc and anti-SrtA antisera. C. Spheroplasts from strain RN6390 producing TspA-Myc were incubated with the indicated concentrations of Proteinase K (pk) at 4°C for 30 mins. A sample of spheroplasts from RN6390 containing pRAB11 (empty) is shown as a negative control. Samples were separated on a 12% bis-tris gel and immunoblotted using anti-Myc, anti-SrtA, anti-EssB and anti-TrxA antisera. D. *S. aureus* RN6390 or the isogenic Δ*essC* or Δ*tspA* strains were cultured in TSB medium and harvested at OD_600_ of 2. Supernatant (sn) and cellular (c) fractions (equivalent of 100 μl culture supernatant and 10 μl of cells adjusted to OD_600_ of 2.0) were separated on bis-Tris gels (15% acrylamide) and immunoblotted using anti-EsxA, EsxC or TrxA antisera. E. Model for organization of TspA in the *S. aureus* envelope. CTD – C-terminal (channel-forming) domain.

TspA is predicted to be 469 amino acids long and to have either one (TMHMM) or two (Predictprotein.org) transmembrane domains towards its C-terminal end. To determine whether it is an integral membrane protein we treated membranes with urea which removes peripherally bound proteins by denaturation. Fig 2B indicates that a large fraction of TspA-Myc was displaced from the membrane to the cytoplasmic fraction by the addition of urea whereas a *bona fide* integral membrane protein, EssB^64-28^, was not displaced by this treatment. We conclude that TspA-Myc peripherally interacts with the membrane. This is consistent with findings from the proteomic experiment as peptides along the entire length of TspA were detected in the secretome (*SI Appendix*, Fig S2).

To determine whether TspA-Myc is exposed at the extracellular side we prepared spheroplasts and treated them with proteinase K. Fig 2C shows that at low concentrations of proteinase K, TspA-Myc was proteolytically cleaved to release a smaller fragment that also cross-reacted with the anti-Myc antibody. At least part of this smaller fragment must be extracellular as it was also degraded as the protease concentration was increased. An approximately 37 kDa C-terminal fragment of TspA-Myc detected natively in the absence of added protease was also extracellular as it was sensitive to digestion by proteinase K. The likely topology of TspA is shown in Fig 2E.

All *S. aureus* T7SS substrate proteins identified to date are found only in a subset of strains, and are linked with specific *essC* subtypes. However, TspA is encoded by all *S. aureus* genomes examined in Warne *et al*.^21^, and is distant from the *ess* locus. This raised the possibility that TspA may be a further secreted core component of the T7 machinery. To examine this, we constructed an in-frame *tspA* deletion in RN6390 and investigated the subcellular location of the T7 secreted component EsxA and the substrate protein EsxC. Fig 2D shows that both EsxA and EsxC are secreted in the absence of TspA. We conclude that TspA is a peripheral membrane protein substrate of the T7SS whose localization and/or stability at the extracellular side of the membrane is dependent on EssC, and that it is not a core component of the T7SS.

### TspA has a toxic C-terminal domain with membrane depolarizing activity that is neutralized by TsaI

Sequence analysis of TspA indicates that homologues are found across the *Staphylococci* (including *S. argenteus, S. epidermidis* and *S. lugdunensis*), in *Listeria* species and Enterococci, but does not provide clues about potential function. However, analysis of TspA using modelling programs predicts strong structural similarity to colicin Ia (Fig 3A), a bacteriocidal protein produced by some strains of *Escherichia coli*. Colicin Ia has an amphipathic domain at its C-terminus that inserts into the cytoplasmic membrane from the extracellular side to form a voltage-gated channel^29-31^. Some limited structural similarity was also predicted with the Type III secretion translocator protein YopB which undergoes conformational changes to forms pores in host cell membranes^32^.

**Figure 3.**
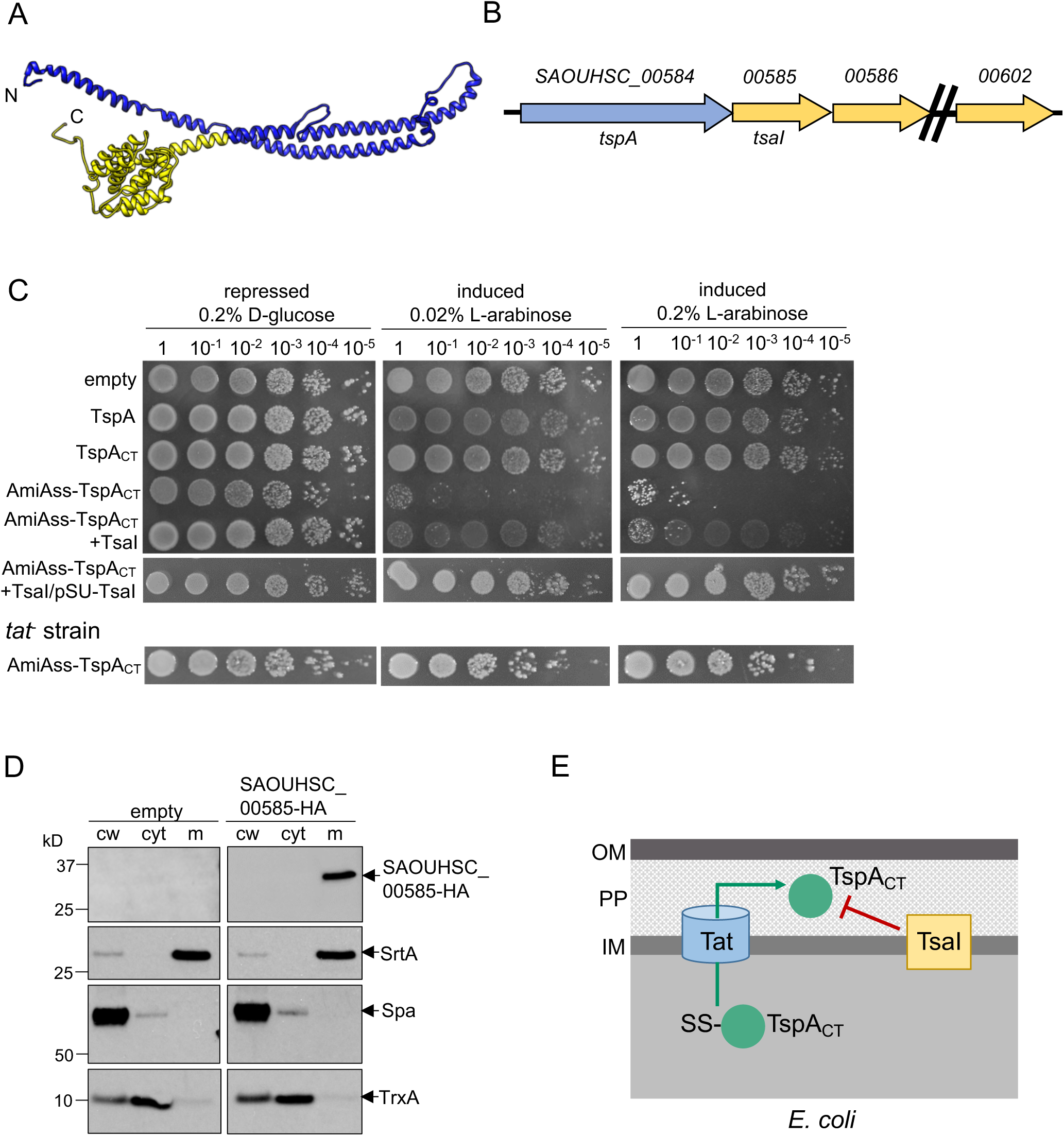
Directed export of TspA C-terminal domain to the periplasm of *E. coli* is toxic. A. Structural model for residues 9-416 of TspA generated using RaptorX (http://raptorx.uchicago.edu/), modelled on the colicin Ia structure ^27^. The predicted channel-forming region is shown in yellow. B. The *tspA* locus. Genes coding for DUF443 proteins are shown in yellow. C. *E. coli* strain MG1655 harboring empty pBAD18-Cm or pBAD18-Cm encoding either full length TspA, the TspA C-terminal domain (TspA_CT_), TspA_CT_ with the AmiA signal sequence at its N-terminus (AmiAss-TspA_CT_), AmiAss-TspA_CT_ and TsaI (SAOUHSC_00585), AmiAss-TspA_CT_/TsaI alongside an additional plasmid-encoded copy of TsaI (from Psu-TsaI) or strain SG3000 (as MG1655, Δ*tatABCD*) harboring pBAD18-AmiAss-TspA_CT_ was serially diluted and spotted on LB plates containing either D-glucose or L-arabinose, as indicated. Plates were incubated at 37°C for 16 hours after which they were photographed. D. *S. aureus* cells harbouring pRAB11 (empty) and pRAB11-SAOUHSC_00585-HA were cultured in TSB medium and expression of SAOUHSC_00585-HA induced by addition of 500 ng/ml ATc when the cells reached OD_600_ of 0.4. The cells were then harvested at OD_600_ of 2. The cells were spun down and subsequently fractionated into cell wall (cw), membrane (m) and cytoplasmic (cyt) fractions. The fractionated samples were separated on bis-tris gels and immunoblotted using the anti-HA antibody, or control anti-sera raised to TrxA (cytoplasmic protein), protein A (SpA, cell wall) or sortase A (SrtA, membrane). E. Schematic representation of the toxicity experiments in Fig 3C, and the inhibition of TspA_CT_ toxicity by the membrane-embedded immunity protein, TsaI. OM, outer membrane; PP, periplasm; IM, inner membrane.

To investigate the function of TspA, DNA encoding full length TspA or the C-terminal domain alone (TspA_CT_) was cloned into a tightly-regulatable vector for expression in *E. coli*. Fig 3C shows that production of TspA or TspA_CT_ did not affect *E. coli* survival. However, colicin Ia shows a sidedness for channel formation because it requires a transmembrane voltage for full insertion^33^. We therefore targeted TspA_CT_ to the periplasm of *E. coli* by fusing to a Tat signal peptide^34, 35^. Fig 3C shows that this construct was toxic, and that toxicity was relieved when the Tat pathway was inactivated (Fig 3C), consistent with the C-terminal domain of TspA exerting toxic activity from the periplasmic side of the membrane.

Bacterially-produced toxins, particularly those that target other bacteria, are often co-expressed with immunity proteins that protect the producing cell from self-intoxication. For example, protection from colicin Ia toxicity is mediated by the membrane-bound Iia immunity protein^36^. TspA is genetically linked to a repeat region of ten genes encoding predicted polytopic membrane proteins with DUF443 domains (Fig 3B). Topological analysis of these proteins predicts the presence of five transmembrane domains with an N_out_-C_in_ configuration. Consistent with this, western blot analysis confirmed that a C-terminally HA-tagged variant of SAOUHSC_00585, which is encoded directly adjacent to TspA, localized to the membrane of *S. aureus* (Fig 3D). To determine whether SAOUHSC_00585 offers protection against the toxicity of the TspA C-terminal domain, we co-produced the AmiAss-TspA_CT_ fusion alongside SAOUHSC_00585 in *E. coli*. Fig 3C shows that co-production of SAOUHSC_00585 offered protection of *E. coli*, particularly when it was constitutively expressed from the pSUPROM plasmid. SAOUHSC_00585 was subsequently renamed TsaI (<u>Ts</u>p<u>A I</u>mmunity protein, Fig 3E).

Pore-forming proteins are widely used as toxins to target either prokaryotic or eukaryotic cells^37, 38^. To assess whether TspA has pore/channel forming activity we investigated whether the production of AmiAss-TspA_CT_ in *E. coli* dissipated the membrane potential. Initially we used the BacLight assay which is based on the dye 3,3′-diethyloxacarbocyanine iodide DiOC_2_(3) that exhibits green florescence in dilute solution but a red shift following membrane potential-driven accumulation in the bacterial cytosol. After sorting of *E. coli* by flow cytometry, the majority of cells harboring the empty vector exhibited red fluorescence, which shifted to green following treatment with the uncoupler carbonyl cyanide 3-chlorophenylhydrazone (CCCP). A similar shift in fluorescence was also observed when *E. coli* produced the AmiAss-TspA_CT_ fusion (Fig 4A), indicative of loss of membrane potential. Co-production of TsaI offered some protection from AmiAss-TspA_CT_-induced depolarization (Fig 4A). We conclude that the C-terminal domain of TspA has membrane depolarizing activity.

**Figure 4.**
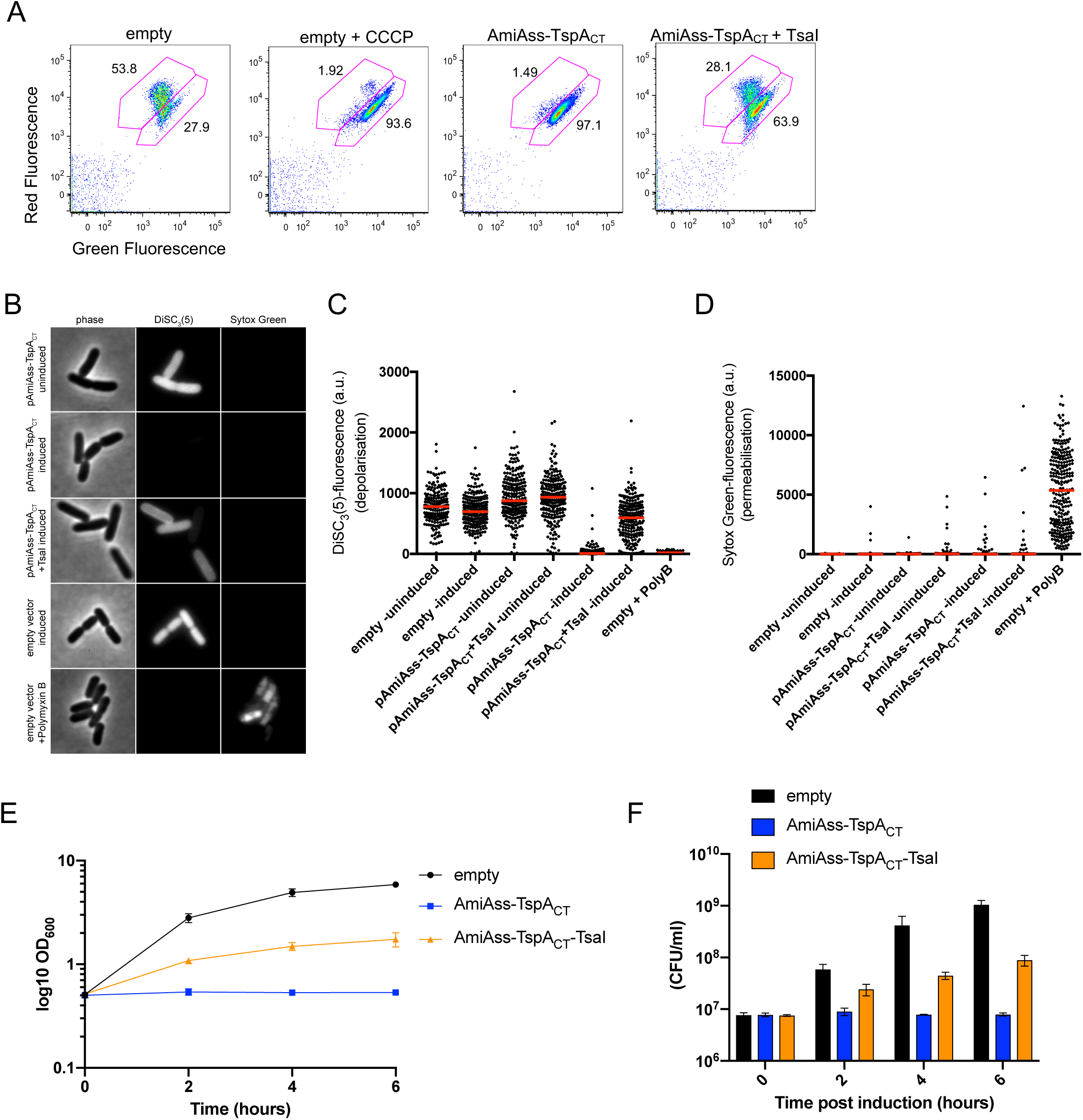
The C-terminal domain of TspA has bacteriostatic activity and disrupts the membrane potential. A. *E. coli* MG1655 harboring pBAD18-Cm (empty), or pBAD18-Cm encoding AmiAss-TspA_CT_ or AmiAss-TspA_CT_ / TsaI were grown in the presence of 0.2% L-arabinose for 1 hour at which point they diluted to 1 × 10^6^ cells per ml and supplemented with 30 μM DiOC_2_(3) for 30 mins. One sample of MG1655 harboring pBAD18 (empty) was also supplemented with 5 μM CCCP at the same time as DiOC_2_(3) addition. Strains were analyzed by flow cytometry. B-D. The same strain and plasmid combinations as A were grown in the presence (induced) or absence (uninduced) of 0.2% L-arabinose for 30 minutes after which they were supplemented with DiSC_3_(5) and Sytox Green and B. imaged by phase contrast and fluorescence microscopy. C+D. Fluorescence intensities of C. DiSC_3_(5) and D. Sytox Green for each sample was quantified using image J. A control sample where Polymyxin B was added to the uninduced empty vector control for 5 minutes before supplemented with DiSC_s_(5) and Sytox Green was included in each experiment. E+F. Growth of *E. coli* MG1655 harboring pBAD18-Cm (empty), or pBAD18-Cm encoding AmiAss-TspA_CT_ or AmiAss-TspA_CT_ / TsaI upon induction with 0.2% L-arabinose. LB medium was inoculated with an overnight culture of *E. coli* strain MG1655 harbouring the indicated constructs to a starting OD_600_ of 0.1. Cells were incubated at 37°C and allowed to reach an OD_600_ of 0.5 (indicated by time 0) before supplementing the growth medium with 0.2% L-arabinose (inducing conditions). The growth was monitored every 2 hours and the colony forming units at each time point was determined (F). Points and bars show mean +/– SEM (*n* = 3 biological replicates).

Membrane depolarization may arise from the formation of ion-selective channels or larger, non-selective pores. To further investigate the mechanism of membrane depolarization we used single-cell microscopy that combines the voltage-sensitive dye DiSC_3_(5) with the membrane-impermeable nucleic acid stain Sytox Green^39^. *E. coli* cells incubated with Polymyxin B, which produces large ion-permeable pores in the *E. coli* cell envelope^40^, showed strong labelling with Sytox Green, indicative of permeabilisation, coupled with very low DiSC_3_(5) fluorescence (Fig 4B-D). By contrast, cells harboring the empty vector had high DiSC_3_(5) fluorescence that was unaffected by supplementation with the inducer L-arabinose, and did not stain with Sytox Green. Cells expressing the AmiAss-TspA_CT_ fusion following incubation with arabinose rapidly depolarized, as evidenced by the marked reduction in DiSC_3_(5) fluorescence, but did not detectably stain with Sytox Green, even after prolonged periods of incubation (Fig 4B-D, *SI Appendix*, Fig S3). Therefore, it appears that TspA acts by triggering membrane depolarization but does so by forming ion channels rather than larger, nonselective pores in the *E. coli* inner membrane. Again, co-production of TsaI significantly protected cells from AmiAss-TspA_CT_-induced depolarization, confirming that it acts as an immunity protein (Fig 4B-D).

Bacterial channel-forming toxins have been reported that have either bacteriocidal^41^ or bacteriostatic^42^ activity. To determine whether the C-terminal domain of TspA was bacteriocidal or bacteriostatic, the growth of *E. coli* producing AmiAss-TspA_CT_ was monitored. It was observed that upon production of AmiAss-TspA_CT,_ *E. coli* ceased to grow (Fig 4E), however quantification of the colony forming units (cfu) indicated that the cells did not lose viability (Fig 4F), pointing to a bacteriostatic action of TspA. We conclude that the C-terminal domain of TspA is a channel-forming toxin with bacteriostatic activity, that is neutralized by the action of TsaI.

### A zebrafish model for *S. aureus* infection and T7SS activity

We next probed whether TspA was important for *S. aureus* virulence, initially through the development of an immunocompetent murine model of *S. aureus* pneumonia. Previous reports have indicated that 2-4 × 10^8^ cfu of strain Newman was a suitable infectious dose, and that bacterial proliferation in lung tissue could be observed after 24 hours^43^. We found that at a dose of 8 × 10^7^ – 2 × 10^8^ of strain RN6390, the mice were asymptomatic and had almost completely cleared the bacteria from lung tissue after 48 hours, whereas a dose of 8 × 10^8^ – 2 × 10^9^ was lethal to all mice within 12 hours. At a dose of 3 × 10^8^, the mice developed symptoms which resolved within 12 hours, and therefore using this dosage we sought to test whether there was a difference in bacterial proliferation and dissemination dependent on the T7SS. However, after 24 hours of infection with 3 × 10^8^ cfu of RN6390 or a cognate Δ*essC* strain, counts recovered from the lungs and livers of mice infected with the Δ*essC* strain were not significantly different than those from mice infected with the wild type (*SI Appendix*, Fig S4).

Given that there are likely to be roles for the T7SS in bacterial competition as well as direct interaction with the host, we next developed a model where these two potentially confounding factors could be investigated. The zebrafish (*Danio rerio*), a widely used vertebrate model for development, has recently been adapted to study bacterial infection by human pathogens^44^. The hindbrain ventricle offers a sterile compartment that can be used to follow bacterial interactions *in vivo*^45^. We first assessed the utility of this infection model by testing the effect of dose and temperature on survival for *S. aureus* inoculated into the hindbrain ventricle of zebrafish larvae 3 days post fertilisation (dpf; *SI Appendix*, Fig S5A). Clear dose-dependent zebrafish mortality was observed, with ∼90% of zebrafish surviving a low dose of *S. aureus* infection (7 × 10^3^ cfu) whereas only ∼55% survived a higher dose (2 × 10^4^ cfu; *SI Appendix*, Fig S5B). Although 28.5°C is the optimum temperature for zebrafish larvae development, *S. aureus* has a temperature optimum of 30 - 37°C for growth. In agreement with this, we observed significantly increased zebrafish mortality at 33°C (relative to 28.5°C) at high dose infection (*SI Appendix*, Fig S5B).

We next assessed whether there was a role for the T7SS in zebrafish mortality. For these experiments, larvae at 3 dpf were inoculated in the hindbrain ventricle with 2 × 10^4^ cfu of RN6390 or an isogenic strain, RN6390 Δ*ess*, lacking all 12 genes (*esxA* through *esaG*) at the *ess* locus^3^, and incubated at 33°C. We routinely observed that zebrafish mortality was significantly reduced, at both 24 and 48 hr post inoculation (hpi), for zebrafish infected with the RN6390 Δ*ess* strain compared to the wild type (*SI Appendix*, Fig S5C; Fig 5A,C). In agreement, total bacterial counts of infected zebrafish revealed that following an initial period of six hours where both strains replicated in a similar manner, there was a significant decrease in recovery of the Δ*ess* strain compared to the wild type after 9 hours (*SI Appendix*, Fig S5D; Fig 5B,D), suggesting that bacteria lacking the T7SS are more rapidly cleared *in vivo*. We also tested a second *S. aureus* strain, COL, in this assay. COL was only weakly virulent at 24 hpi, but at high dose substantial mortality was seen after 48 hr (*SI Appendix*, Fig S6A). As before, zebrafish mortality was at least in part dependent on a functional T7SS (*SI Appendix*, Fig S6B), although we observed no difference in bacterial burden between the wild type and Δ*essC* strain at the timepoints sampled (*SI Appendix*, Fig S6C). We conclude that the T7SS plays a role in virulence of *S. aureus* in this zebrafish infection model.

**Figure 5.**
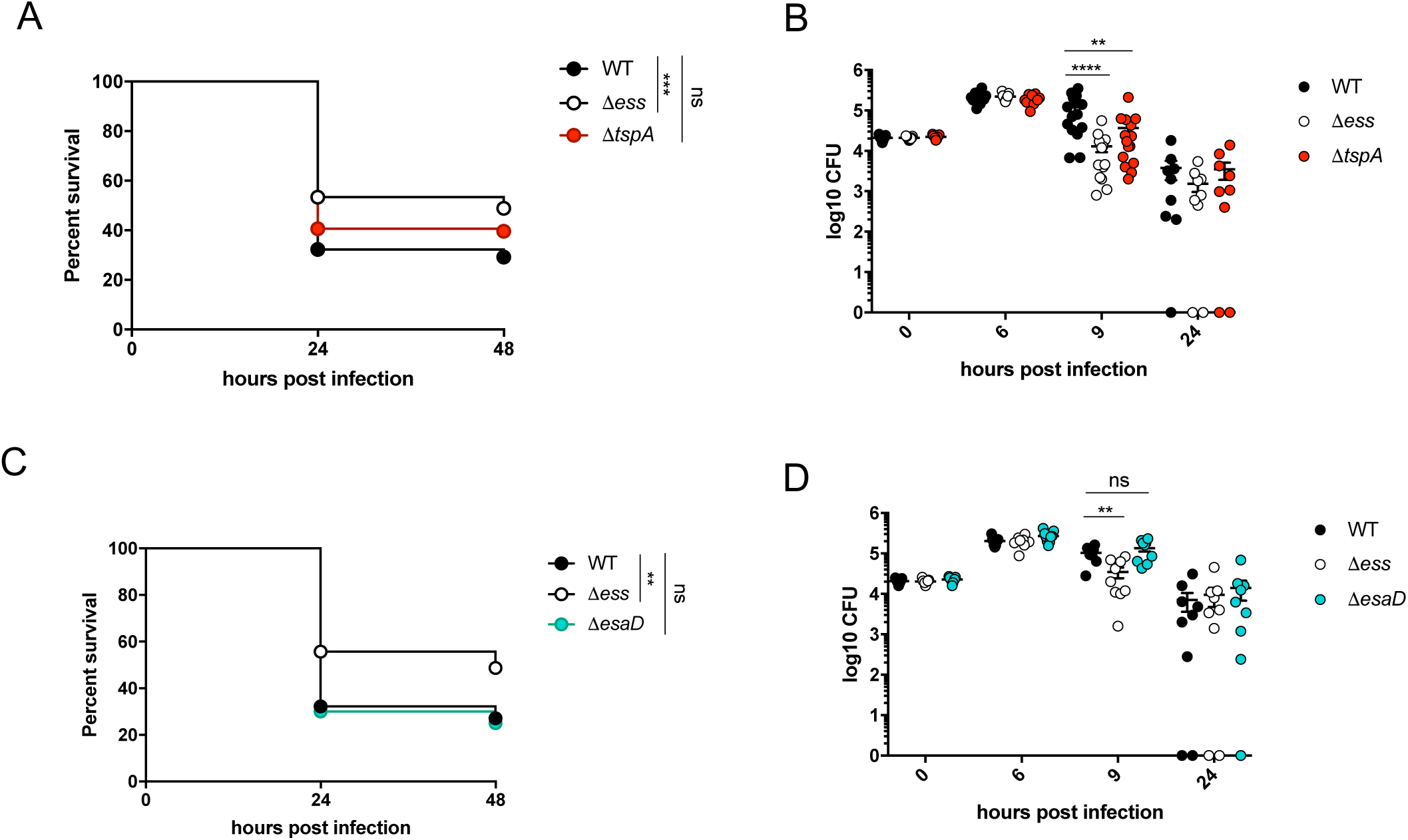
The T7SS contributes to virulence in a zebrafish infection model. A. Survival curves of 3 dpf zebrafish *lyz*:dsRed larvae infected in the hindbrain ventricle with RN6390-gfp (WT) or otherwise isogenic Δ*ess*-gfp or Δ*tspA*-gfp strains at a dose of ∼2 × 10^4^ cfu and incubated at 33°C for 48 hpi. Data are pooled from four independent experiments (*n*=25-51 larvae per experiment). Results were plotted as a Kaplan-Meier survival curve and the *p* value between conditions was determined by log-rank Mantel-Cox test. B. Enumeration of recovered bacteria at 0, 6, 9 or 24 hpi from zebrafish larvae infected with the same strains as A. Pooled data from 3 independent experiments. Circles represent individual larva, and only larvae that survived the infection were included. No significant differences observed between strains at 0, 6 or 24 hpi. Mean ± SEM also shown (horizontal bars). Significance testing was performed using a one-way ANOVA with Sidak’s correction at each timepoint. C. Survival curves of 3 dpf zebrafish *lyz*:dsRed larvae infected in the hindbrain ventricle with RN6390-gfp (WT) or otherwise isogenic Δ*ess*-gfp, Δ*esaD*-gfp strains at a dose of ∼2 × 10^4^ cfu and incubated at 33°C for 48 hpi. Data are pooled from three independent experiments (*n*=26-32 larvae per experiment). Results are plotted as a Kaplan-Meier survival curve and the *p* value between conditions was determined by log-rank Mantel-Cox test. D. Enumeration of recovered bacteria at 0, 6, 9 or 24 hpi from zebrafish larvae infected with the strains as C. Pooled data from 3 independent experiments. Circles represent individual larva, and only larvae having survived the infection were included. No significant differences observed between strains at 0, 6 or 24 hpi. Mean ± SEM also shown (horizontal bars). Significance testing was performed using a one-way ANOVA with Sidak’s correction at each timepoint. \*\**p*<0.01, *** *p*<0.001, **** *p*<0.0001, ns, not significant.

In addition to TspA, a second T7SS secreted toxin, EsaD (also called EssD^9, 46^), a nuclease, has been identified in some *S. aureus* strains. EsaD was shown to inhibit growth of a competitor *S. aureus* strain *in vitro*^6^, but has also been directly implicated in virulence through modulation of cytokine responses and abscess formation^9, 46^. We therefore determined whether TspA or EsaD was required for virulence in the zebrafish infection model. Infection of larvae with strain RN6390 lacking TspA resulted in levels of mortality intermediate between the wild type and Δ*ess s*train (Fig 5A), and a significantly reduced bacterial burden relative to the wild type strain at 9 hpi (Fig 5B). By contrast, no difference was observed in either zebrafish mortality (Fig 5C) or bacterial burden (Fig 5D) between infection with RN6390 and an isogenic *esaD* mutant, indicating no detectable role of EsaD in virulence. Taken together, we conclude that zebrafish infection can be used to investigate the role of T7SS effectors *in vivo*, and that TspA (but not EsaD) contributes to T7SS-mediated bacterial replication *in vivo*.

Previous studies have shown that the T7SS of *S. aureus* is involved in modulating the murine host immune response^9, 46^. To test whether altered immune responses mediate the increased clearance of the Δ*ess* and Δ*tspA* deletion strains at 9 hpi, we investigated the role of the T7SS in the zebrafish larval cytokine response during *S. aureus* infection *in vivo* (*SI Appendix*, Fig S7). The expression of two host pro-inflammatory markers interleukin 8 (*cxcl8)* and interleukin 1 beta (*il-1b*) were quantified using qRT-PCR in larvae infected with 2 × 10^4^ cfu of RN6390 wild type, Δ*ess*, Δ*tspA* and Δ*esaD* strains. In comparison to PBS injected larvae, *S. aureus* infection caused a robust increase in both *cxcl8* and *il-1b* expression at 6 hpi (when the bacterial burden among strains was similar; *SI Appendix*, Fig S7). However, no significant difference in gene expression was observed among larvae infected with wild type and any of the three deletion strains (Δ*ess*, Δ*tspA* and Δ*esaD*; *SI Appendix*, Fig S7).

Neutrophils represent the first line of defence against *S. aureus* infection^47^ and the recently discovered substrate of EssC variant 2 strains, named EsxX, has been implicated in neutrophil lysis, therefore contributing to evasion of the host immune system^48^. By contrast, the T7SS of *Mycobacterium tuberculosis* (ESX-1) is associated with manipulation of the inflammatory response during infection allowing for bacterial replication in macrophages^49-52^. To investigate whether the *S. aureus* T7SS modulates interaction with leukocytes, we analysed the recruitment of immune cells to the hindbrain using two transgenic lines in which dsRed is expressed specifically in neutrophils (Tg(*lyz*::dsRed)) or mCherry is expressed specifically in macrophages (Tg(*mpeg*::Gal4-FF)^gl25^/Tg(*UAS*-*E1b*::*nfsB*.mCherry)^c264^, herein Tg(*mpeg1*::G/U::mCherry)). Zebrafish larvae were infected with RN6390 wild type, Δ*ess* and Δ*tspA* strains in the hindbrain ventricle at 3 dpf and imaged under a fluorescent stereomicroscope at 0, 3 and 6 hpi in order to monitor neutrophil (*SI Appendix*, Fig S8A+B) and macrophage (*SI Appendix*, Fig S8C+D) behavior. In zebrafish larvae infected with *S. aureus*, a significant increase in neutrophil recruitment to the hindbrain ventricle was detected in comparison to PBS-injected larvae at both 3 and 6 hpi (*SI Appendix*, Fig S8B). However, no difference in neutrophil recruitment to the Δ*ess* and Δ*tspA* strains relative to wild type was detected at any of the time points tested (*SI Appendix*, Fig S8B). Similar to the neutrophil recruitment experiments, a significant increase in macrophage recruitment to the site of *S. aureus* infection was observed when compared to PBS injected larvae at 3 hpi (*SI Appendix*, Fig S8D). However, there was no significant difference in macrophage recruitment among the wild type and T7SS mutant strains (*SI Appendix*, Fig S8D).

### T7SS dependent bacterial competition *in vivo*

Although TspA is required for optimal *S. aureus* virulence in the zebrafish model, the observed toxicity when heterologously produced in *E. coli* coupled with the presence of immunity genes encoded downstream of *tspA* strongly suggested that secreted TspA may also have antibacterial activity. Previously, to observe antibacterial activity of the nuclease EsaD in laboratory growth media required the toxin to be overproduced from a multicopy plasmid^6^. However, zebrafish larvae have recently been adapted to study bacterial predator-prey interactions^45^, and we reasoned that since the T7SS was active in our zebrafish infection model it may also provide a suitable experimental system to investigate the impact of T7-mediated bacterial competition *in vivo*.

In these experiments we used COL as the attacker strain and RN6360 and its derivatives as the target; it should be noted that these strains produce the same TspA and EsaD isoforms as well as similar suites of immunity proteins. COL was co-inoculated into the hindbrain ventricle, at a 1:1 ratio, with either RN6390 or an isogenic strain lacking all potential immunity proteins for EsaD and TspA (FRU1; RN6390 Δ*saouhsc00268-00278*, Δ*saouhsc00585-00602*). A significant reduction in recovery of the target strain lacking immunity genes was observed compared to the isogenic parental strain at 15 hr post infection (Fig 6A). Conversely, there was significantly greater zebrafish mortality at 24 hr after co-inoculation of COL with the wild type RN6390 than the immunity mutant strain (Fig 6B). Since COL is almost completely avirulent at this time-point (*SI Appendix*, Fig S6) we infer that mortality arises from RN6390, and as the wild type strain survives better than the immunity deletion strain when co-inoculated with COL, this accounts for the greater zebrafish mortality.

**Figure 6.**
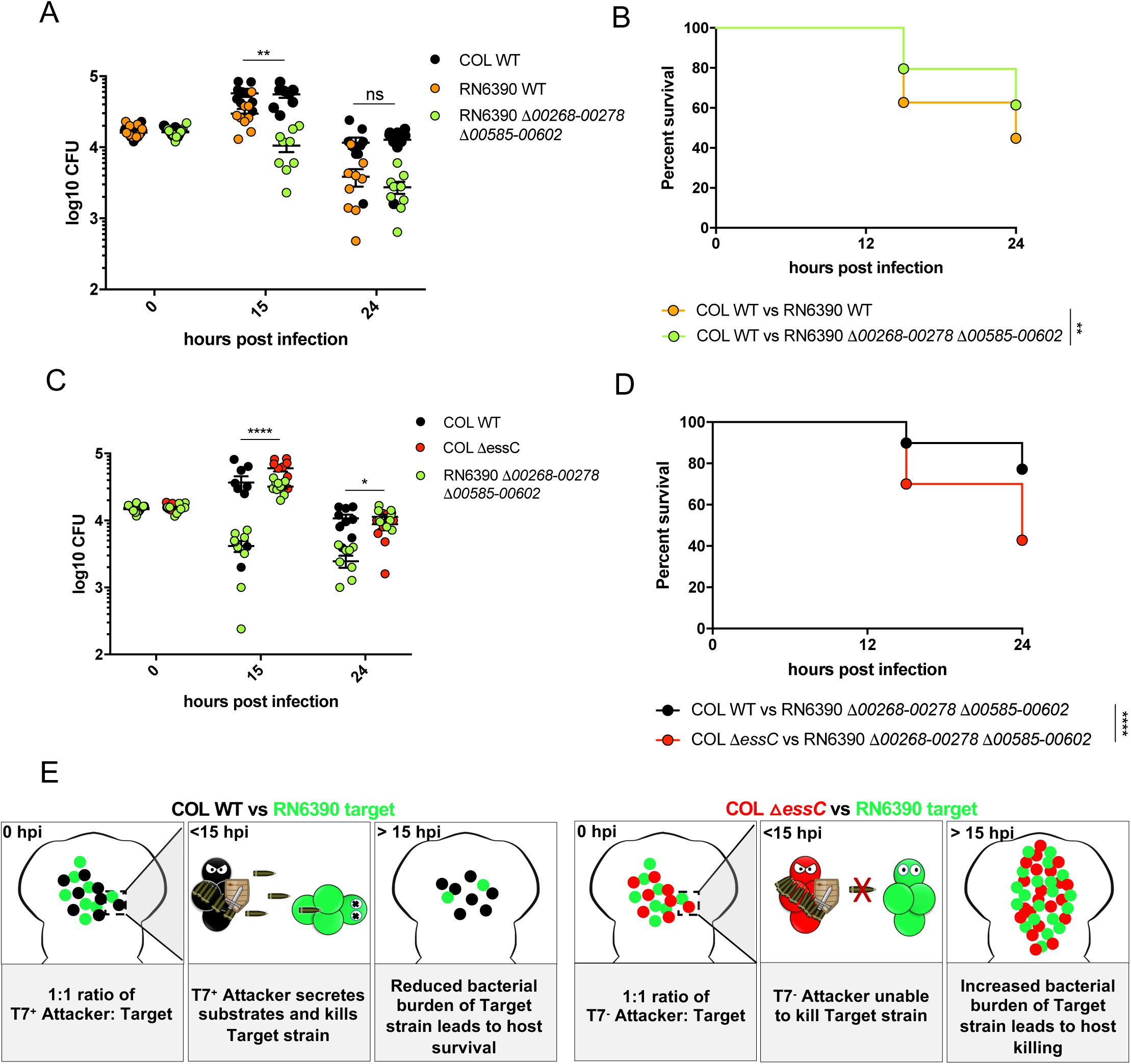
Development of an *in vivo* model to study bacterial competition. Wild type AB zebrafish larvae at 3 dpf were co-infected with a 1:1 mix of an attacker strain (either COL-mCherry (WT) or COL Δ*essC*-mCherry as indicated) and a target strain (either RN6390-gfp (WT) or RN6390 Δ*00268-278* Δ*00585-00602*-gfp, as indicated). A+C. Enumeration of recovered attacker and prey bacteria from zebrafish larvae at 0, 15 or 24 hpi. Pooled data from 3 independent experiments, Mean ± SEM also shown (horizontal bars). Significance testing performed by unpaired *t* test. B+D. Survival curves of zebrafish injected with the indicated strain pairs. Data are pooled from three independent experiments. Results are plotted as a Kaplan-Meier survival curve and the p value between conditions was determined by log-rank Mantel-Cox test. \*\**p*<0.01, *** *p*<0.001, **** *p*<0.0001, ns, not significant. E. Model highlighting the role for the T7SS in competition *in vivo*.

To confirm that reduced growth of the RN6390 immunity mutant strain was dependent upon a functional T7SS in the attacking strain, we repeated the co-inoculation experiments using a T7 mutant strain of COL (COL Δ*essC*). The RN6390 immunity mutant strain showed significantly higher recovery after 15 hours in the presence of the COL T7 mutant strain than wild type COL (Fig 6C) and accordingly this was linked with reduced zebrafish survival (Fig 6D). Collectively, these data highlight the utility of zebrafish for investigating *S. aureus* competition *in vivo*, and demonstrate that bacterial competition and zebrafish mortality is dependent on a functional T7SS in the attacking strain (COL). This is outlined in the schematic (Fig 6E). Conversely, the ability of the prey strain (RN6390) to survive T7-dependent killing is dependent upon the immunity proteins against EsaD and TspA, because when these are not present, fewer bacteria are recovered.

Finally, we investigated which of the EsaD and TspA toxins was responsible for inter-strain competition by using variants of COL deleted for either *tspA* or *esaD* as the attacking strain. It was seen that in the absence of either TspA (Fig 7A) or EsaD (Fig 7C) there was a significant increase in recovery of the RN6390 Δ*saouhsc00268-00278*, Δ*saouhsc00585-00602* prey strain, indicating that each of these toxins has activity against the target strain. However, there was a more pronounced increase in zebrafish mortality when the attacker strain lacked *esaD* than *tspA* (compare Figs 7B and D), suggesting that EsaD has the more potent antibacterial activity in these conditions.

**Figure 7.**
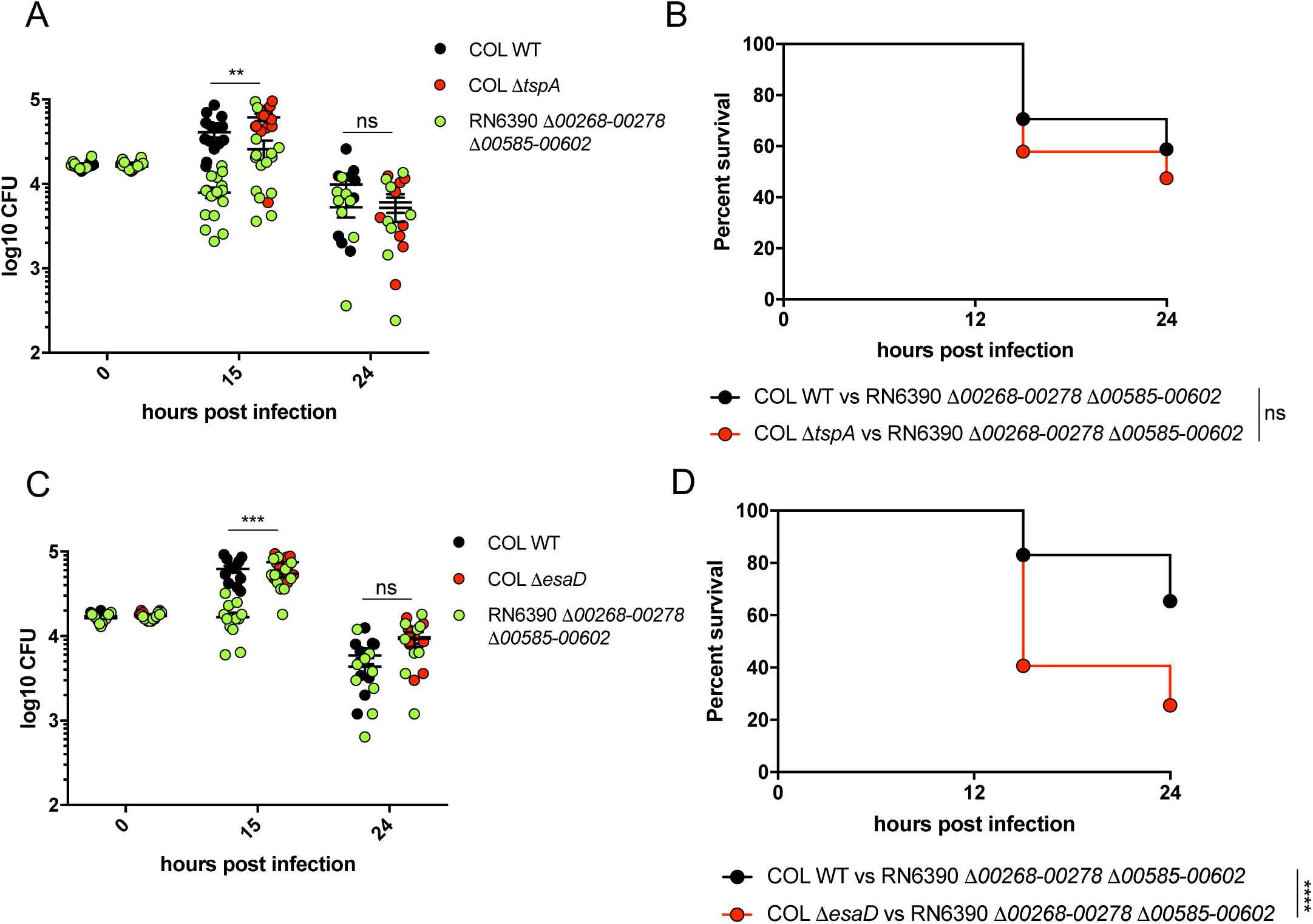
TspA and EsaD dependent bacterial competition *in vivo*. Wild type AB zebrafish larvae at 3 dpf were co-infected with a 1:1 mix of an attacker strain (either COL-mCherry (WT), COL Δ*tspA*-mCherry or COL Δ*esaD*-mCherry as indicated) and a target strain (RN6390 Δ*00268-278* Δ*00585-00602*-gfp). A+C. Enumeration of recovered attacker and prey bacteria from zebrafish larvae at 0, 15 or 24 hpi. Pooled data from 3 independent experiments, Mean ± SEM also shown (horizontal bars). Significance testing performed by unpaired *t* test. B +D. Survival curves of zebrafish injected with the indicated strain pairs. Data are pooled from three independent experiments. Results are plotted as a Kaplan-Meier survival curve and the *p* value between conditions was determined by log-rank Mantel-Cox test. \*\**p*<0.01, *** *p*<0.001, **** *p*<0.0001, ns, not significant.

## Discussion

Here we have taken an unbiased approach to discover substrates of the T7SS in *S. aureus* RN6390, identifying the LXG-domain protein, TspA. TspA localizes to the cell envelope and has a toxic C-terminal domain that has membrane-depolarizing activity. While all other previously-identified T7 substrates are encoded at the *ess* locus and are associated with specific *essC* subtypes^21, 48^, TspA is encoded elsewhere on the genome, and is conserved across all *S. aureus* strains. This suggests TspA plays a key role in *S. aureus*, and indeed we show using a zebrafish infection model that it contributes to T7SS-mediated bacterial replication *in vivo*.

Pore- and channel-forming toxins are key virulence factors produced by many pathogenic bacteria^53^, that can act both extracellularly to form pores in eukaryotic cells, like some bacterial hemolysins^54^, or intracellularly for example by altering permeability of the phagosome, like the pore-forming toxin Listeriolysin-O, or the Type III secretion system effector VopQ^55, 56^. It should be noted that the T7SS of strain USA300 has been shown to play a role in release of *S. aureus* during intracellular infection^57^, however RN6390 is only poorly invasive in bronchial epithelial cell lines, and intracellular survival and bacterial release is independent of the T7SS (data not shown). The *S. aureus* T7SS has been strongly linked with modulating the host innate immune response^9, 45^. However, we did not observe any significant difference between wild type and T7SS mutant strains in modulating cytokine expression and phagocyte recruitment in zebrafish larvae. Although the precise mechanism by which the T7SS and TspA interacts with host cells remains to be determined, we hypothesise that the T7SS plays a role after phagocytosis by immune cells to influence intracellular survival. Future work using high resolution single cell microscopy would allow for individual *S. aureus* cells, as well as their interactions with neutrophils and/or macrophages, to be tracked *in vivo*.

Sequence alignments indicate that the C-terminal domain of TspA is polymorphic across *S. aureus* strains (Fig S9) and structural modelling of TspA suggests homology to colicin Ia. Colicin Ia is a toxin that forms voltage-gated ion channels in the plasma membrane of sensitive *E. coli* strains. The formation of these channels results in lysis of target bacteria^30, 41^. Heterologous expression of the C-terminal predicted channel-forming domain of TspA was shown to dissipate the membrane potential of *E. coli* when it was targeted to the periplasm, probably through formation of an ion channel. Unlike Colicin Ia, however, heterologous production of the TspA toxin domain was associated with a bacteriostatic rather than a bacteriocidal activity. Colicins and pyocins are also examples of polymorphic toxins^38^ and the producing cells are generally protected from colicin-mediated killing by the presence of immunity proteins^36^. A cluster of membrane proteins from the DUF443 domain family are encoded downstream of *tspA*, and we show that at least one of these (SAOUHSC_00585; TsaI) acts as an immunity protein to TspA by protecting *E. coli* from TspA-induced membrane potential depletion.

Polymorphic toxins are frequently deployed to attack competitor bacteria in polymicrobial communities^37^, and there is growing evidence that a key role of the T7SS in some bacteria is to mediate inter- and intra-species competition^6,7^. In addition to TspA, many commonly-studied strains of *S. aureus*, including RN6390 and COL also secrete a nuclease toxin, EsaD^6^. We adapted our zebrafish larval infection model to assess the role of the T7SS and the secreted toxins TspA and EsaD in intra-species competition. We observed that strain COL was able to outcompete RN6390 in a T7SS-dependent manner in these experiments, provided that RN6390 was lacking immunity proteins to TspA and EsaD. Experiments with individual COL attacker strains deleted for either *tspA* or *esaD* showed that each of the toxins contributed to the competitiveness of COL in these assays. As *S. aureus* is a natural coloniser of human nares and can also exist in polymicrobial communities in the lungs of cystic fibrosis patients, we suggest that secreted T7 toxins including TspA allow *S. aureus* to establish its niche by outcompeting other bacteria. Indeed, the observation that the T7SS gene cluster is highly upregulated in the airways of a cystic fibrosis patient^5^ would be consistent with this hypothesis.

LXG domain proteins appear to form a large substrate family of the firmicutes T7SS. Three LXG domain proteins of *Streptococcus intermedius* have been shown to mediate contact-dependent inhibition^7^, and the association of TspA with the *S. aureus* cell envelope would also imply that toxicity is contact-dependent. The LXG domain is predicted to form an extended helical hairpin, which could potentially span the cell wall, displaying the toxin domain close to the surface. How any of these toxin domains reach their targets in the prey cell is not clear. One possibility is that the toxin domain is taken up into the target cell upon interaction with a surface receptor, as observed for the Type V-dependent contact inhibition systems in Gram-negative bacteria^58, 59^. During this process the CdiA protein, which also has a C-terminal toxin domain, is proteolyzed, releasing the toxin to interact with its cellular target^60^. Further work would be required to decipher the mechanism/s by which LXG toxins access target cells and whether the toxin domains undergo proteolysis to facilitate cellular entry.

In conclusion, channel forming toxin substrates have been associated with other protein secretion systems^42, 55, 56, 58^, but this is the first to be functionally described for the T7SS. To our knowledge it is only the second bacterial exotoxin identified to have a phenotype in both bacterial competition and virulence assays, after VasX from *Vibrio cholerae*^61, 62^.

## Materials and Methods

### Bacterial strains, plasmids and growth conditions

Construction of strains and plasmids, and growth conditions are described in *SI Materials and Methods*. Plasmids and strains used in this study are given in *SI Appendix*, Tables S2 and S3.

### Mass spectrometry data analysis and label-free quantitation

Preparation of *S. aureus* culture supernatants for proteomic analysis is detailed in *SI Materials and Methods*. Sample preparation and mass spectrometry analysis was performed similar to previously described work^63-66^ and detailed methods are described in *SI Materials and Methods*.

### Cell fractionation and western blotting

Preparation of *S. aureus* cell and supernatant samples for western blotting, and subcellular fractionation of *S. aureus* into cell wall, membrane and cytoplasmic fractions were as described previously^3^. Preparation of urea-washed membrane fractions was adapted from Keller *et al*.^67^. Briefly, broken cell suspensions were thoroughly mixed with a final concentration of 4M urea and incubated for 20 minutes at room temperature. Membranes were harvested by ultracentrifugation (227 000 x *g*, 30 min). The supernatant was retained as the urea-treated cytoplasmic fraction and the membrane pellet resuspended in 1 x PBS, 0.5 % Triton X-100. For spheroplast preparation, the method of Gotz *et al*.^68^ was adapted. Briefly, strains were cultured as described above, cells were harvested at OD_600_ of 2.0 and resuspended in Buffer A (0.7M sucrose, 20mM maleate, 20mM MgCl_2_, pH 6.5). Lysostaphin and lysozyme were added at 20 μg/ml and 2 mg/ml final concentration, respectively, and cells incubated at 37°C for 1 hour. Cell debris was pelleted by centrifugation (2,500 x *g* for 8 min) and the resulting supernatant centrifuged at 16,000 x *g* for 10 min to pellet the spheroplasts. Spheroplasts were resuspended in Buffer A and treated with increasing concentrations of Proteinase K on ice for 30 mins. 0.5 mM phenylmethylsulfonyl fluoride was added to terminate the reaction and samples mixed with 4x Nu PAGE LDS sample buffer and boiled for 10 min prior to further analysis. Western blotting was performed according to standard protocols using the following antibody dilutions α-EsxA 3 1:2500; α-EsxC^3^ 1:2000, α-EssB^3^ 1:10000, α-TrxA^69^ 1:25000, α-SrtA (Abcam, catalogue number ab13959) 1:3000, α-HA (HRP-conjugate, Sigma catalogue number H6533) α-Myc (HRP-conjugate, Invitrogen catalogue number R951-25) 1:5000, and goat anti Rabbit IgG HRP conjugate (Bio-Rad, catalogue number 170-6515) 1:10000.

### Bacterial membrane potential detection

To assess bacterial membrane potential, the method of Miyata *et al*.^70^ was adapted, using the BacLight bacterial membrane potential kit (Invitrogen). Detailed methods to assess both bacterial membrane potential and permeabilization are described in *SI Materials and Methods*.

### Zebrafish Infection

Wild-type (WT, AB strain) or transgenic Tg(*lyz*::dsRed)^*nz50*^ 71 zebrafish were used for all experiments. Embryos were obtained from naturally spawning zebrafish, and maintained at 28.5°C until 3 days post fertilisation (dpf) in embryo medium (0.5x E2 medium supplemented with 0.3 g/ml methylene blue)^72^. For injections, larvae were anesthetized with 200 µg/ml tricaine (Sigma-Aldrich) in embryo medium. Hindbrain ventricle infections were carried out at 3 dpf and incubated at 33°C unless specified otherwise. Bacteria were subcultured following overnight growth until they reached OD_600_ of 0.6. For injection of larvae, bacteria were recovered by centrifugation, washed and resuspended in 1x PBS, 0.1% phenol red, 1% polyvinylpyrrolidone to the required cfu/ml. Anaesthetized larvae were microinjected in the hindbrain ventricle (HBV) with 1–2 nL of bacterial suspension. At the indicated times, larvae were sacrificed in tricaine, lysed with 200 μL of 0.4% Triton X-100 and homogenized mechanically. Larval homogenates were serially diluted and plated onto TSB agar. Only larvae having survived the infection were included for enumeration of colony forming units (cfu). For zebrafish virulence assays all *S. aureus* strains were chromosomally tagged with GFP which included RN6390 wild type, and isogenic Δ*ess*, Δ*tspA* and Δ*esaD* strains. For *in vivo* competition experiments, COL (attacker) strains were chromosomally-tagged with mCherry and RN6390 (target) strains with GFP. Attacker and target strains were subcultured, harvested and resuspended in PBS as above. Attacker and target strains were mixed at a 1:1 ratio and injected in the hindbrain ventricle, with 1-2 nL of bacterial suspension. Larvae were sacrificed at 15 hpi or 24 hpi, serially diluted and plated on TSB agar, and attacker and target strains were enumerated by fluorescence (GFP and mCherry). Quantitative reverse transcription PCR and *S. aureus* - leukocyte microscopy methods are described in *SI Materials and Methods*.

## Acknowledgements

This study was supported the Wellcome Trust [through Early Postdoctoral Training Fellowship for Clinician Scientists WT099084MA to JDC, Investigator Award 110183/Z/15/Z to TP and Institutional Strategic Support Fund 105606/Z/14/Z to the University of Dundee], the UK Biotechnology and Biological Sciences Research Council [grants BB/H007571/1 and EASTBIO DTP1 grant ref BB/J01446X/1], The Microbiology Society (through a Research Visit Grant awarded to FRU) and a China Scholarship Council PhD studentship (to ZC). MT was funded by the Medical Research Council UK through grant MC_UU_12016/5. Work in the SM laboratory is supported by a European Research Council Consolidator Grant (772853 - ENTRAPMENT), Wellcome Trust Senior Research Fellowship (206444/Z/17/Z) and the Lister Institute of Preventive Medicine. We thank Professor Frank Sargent and Dr Sabine Grahl for providing us with *E. coli* strain SG3000, Professor Jan-Maarten van Dijl (University of Groningen) for the kind gift of anti-TrxA antiserum, and Drs Giuseppina Mariano, Sarah Coulthurst, Vincenzo Torraca, and Melanie Blokesch for helpful discussion and advice.

## SI Materials and Methods

### Bacterial strains, plasmids and growth conditions

All strains and plasmids used in this study are given in *SI Appendix*, Tables S2 and S3. *S. aureus* strain RN6390^1^ and its Δ*essC*^2^, Δ*esaD*^3^, Δ*SAOUHSC_00268-00278*^3^ and Δ*ess* (Δ*esxA*-*esaG*)^2^ derivatives along with strain COL^4^ have been described previously. An in-frame deletion of *tspA* (*SAOUHSC_00584*) in RN6390 was constructed by allelic exchange using plasmid pIMAY (*SI Appendix*, Table S3)^4^. The upstream and downstream regions including the start codon and last six codons were amplified from RN6390 genomic DNA using primers listed in *SI Appendix*, Table S4 and were cloned into pIMAY and introduced onto the chromosome by recombination as described previously^4^. For deletion of *tsaI* and its homologues, the upstream regions of *SAOUHSC_00585* including its first four codons and the downstream *SAOUHSC_00602* including its last four codons were amplified from RN6390 genomic DNA, cloned into pIMAY and was introduced into strain RN6390 Δ*SAOUHSC_00268-00278* to generate strain FRU1 (as RN6390 Δ*SAOUHSC_00268-00278*, Δ*SAOUHSC_00585-00602*). For in frame deletion of *esaD* (SACOL0281) and *tspA* (SACOL0643) in strain COL, constructs pIMAY-esaD^3^ and pIMAY-tspA were used, following the protocol of Monk *et al*.^5^. Derivatives of strains harboring markerless *gfp* or *mCherry* chromosomal insertions were constructed according to de Jong *et al*.^6^ using plasmids pTH100 and pRN111, respectively. *E. coli* strain JM110 was used for cloning purposes and MG1655^7^ and its isogenic Δ*tatABCD* derivative SG3000^8^ was used for toxicity assays.

All oligonucleotides used in this study are listed in *SI Appendix*, Table S4, and RN6390 chromosomal DNA was used as template unless otherwise stated. Plasmid pRAB11-tspA-myc encodes TspA with a C-terminal Myc tag in pRAB11^9^ and was constructed following amplification with primers tspA cmyc fw and tspA cmyc rv. Plasmid pRAB11-02448-ha produces SAOUHSC_02448 with a C-terminal HA tag from vector pRAB11 and the encoding gene was amplified using primers 02448 cha fw and 02448 cha rv. Plasmid pRAB11-00389-ha codes for SAOUHSC_00389 with a C-terminal HA tag in pRAB11 and was constructed following amplification with primers 00389 cha fw and 00389 cha rv. Plasmid pRAB11-00406-myc encodes SAOUHSC_00406 with a C-terminal Myc tag in pRAB11 and was constructed following amplification with primers 00406 cmyc fw and 00406 cmyc rv. In each case the amplified gene is preceded by the *esxA* RBS (AGGAGGTTTCTAGTT), and were cloned as *Kpn*I - *Sac*I fragments. Plasmid pRAB11-00585-myc encodes SAOUHSC_00585 with a C-terminal HA tag in pRAB11 and was constructed following amplification with primers 00585 cha fw and 00585 cha rv, digestion with *Bgl*II and *Eco*RI and cloning into similarly cut pRAB11. Plasmid pRMC2-ssaA-ha codes for SsaA with a C-terminal HA tag. It was constructed following amplification of SsaA using primers pRMC2-ssaA-bglII-for and pRMC2-ssaA-HA-rev-EcoRI, digestion with *Bgl*II and *Eco*RI and cloning into similarly cut pRMC2^10^.

Plasmid pBAD18-tspA codes for the full length TspA. The encoding gene was amplified using primers tspA fl fw and tspA fl rv, digested with *Xba*I and *Sal*I and subsequently cloned into similarly cut pBAD18-Cm^11^. Plasmid pBAD18-tspA_CT_ encodes for the last 251aa of TspA; the encoding DNA was amplified using primers tspA cp fw and tspA fl rv, digested with *Xba*I and *Sal*I and cloned into similarly cut pBAD18-Cm. Plasmid pBAD18-AmiAss-tspA_CT_ codes for the *E. coli* AmiA signal sequence fused in-frame to the N-terminus of TspA_CT_. This was constructed following separate amplification of the DNA encoding the first 36 amino acids of AmiA using primers AmiAss fw and AmiAss rv with *E. coli* MG1655 chromosomal DNA as template digestion with *Nhe*I and *Kpn*I and cloning into similarly cut pBAD18-Cm. The TspA_CT_ coding sequence was then amplified using tspA cp fw (which lacks a start codon) and tspA fl rv, digested with *Xba*I and *Sal*I and cloned into similarly cut pBAD18-AmiAss. Plasmid pBAD18-AmiAss-tspA_CT_ + tsaI codes for the AmiAss-TspA_CT_ fusion along with TsaI. The *tsaI* (*saouhsc_00585*) gene was amplified using primers tsaI fw and tsaI rv, digested with *Sal*I and *Sph*I and cloned into similarly cut pBAD18-AmiAss-tspA_CT_.

Plasmid pSUPROM-TsaI produces TsaI constitutively from the *E. coli tat* promoter. The encoding gene was amplified using primers pSU-TsaI fw and pSU-TsaI rv, digested with *Bam*HI and *Xho*I and cloned into similarly-digested pSUPROM^12^. *S. aureus* strains were cultured in RPMI medium for proteomic analysis, as detailed below. For all other experiments *S. aureus* strains were grown in TSB medium at 37°C with vigorous agitation. Chloramphenicol was used 10 µg/ml final concentration for plasmid selection. Anhydrotetracycline (ATC) was added to *S. aureus* cultures at 1 µg/ml during allelic gene replacement. For induction of plasmid-encoded proteins, 500 ng/ml ATC was added to cultures at OD_600_ of 0.4, and cells were harvested at OD_600_ of 2.0. *E. coli* was grown aerobically in LB at 37°C, supplemented with ampicillin (100 µg/ml), kanamycin (50 µg/ml) or chloramphenicol (25 µg/ml) where appropriate. D-glucose and L-arabinose were used to control expression of cloned genes from the pBAD18-Cm vector^11^. For toxicity assays single colonies of MG1655 freshly transformed with the appropriate pBAD18-Cm construct growing on LB agar containing 0.2% D-glucose were picked and re-suspended in LB to an OD_600_ of 1.0. Serial dilutions to 10^−5^ were prepared and 5 μL of each spotted onto LB agar supplemented with 0.2% D-glucose, 0.02% L-arabinose or 0.2% L-arabinose and incubated overnight at 37°C. For growth curve measurements, overnight cultures of MG1655 harboring pBAD18-Cm constructs were sub-cultured to a starting OD_600_ of 0.1 (t=0) and incubated at 37°C and allowed to reach an OD_600_ of 0.5 before supplementation with 0.2% L-arabinose. Optical density readings at 600nm were taken for a 6 hour growth period, with readings collected every 2 hours. The number colony forming units was calculated at t=0 and every 2 hours post-induction with L-arabinose. Serial dilutions to 10^−6^ were prepared and 100 μL of each plated onto LB agar containing 25 µg/ml chloramphenicol and incubated overnight at 37°C.

### Preparation of culture supernatants for proteomic analysis

*S. aureus* strains were grown overnight in 2 mL TSB after which cells were harvested, washed three times with 10 mL of RPMI medium, resuspended in 2 mL of RPMI and used to inoculate 200 mL RPMI in 2 L baffled flasks. Cultures were grown at 37°C with vigorous agitation until an OD_600_ of 1.0 was reached, after which cultures were cooled to 4°C, cells pelleted and supernatant proteins precipitated with 6% trichloroacetic acid (TCA) on ice overnight. The precipitated protein samples were harvested by centrifugation (15 min at 18000 *g*) re-suspended in 80% acetone (−20°C) and washed twice with 80% acetone. Pellets were air dried at room temperature and transferred to the mass-spectrometry facility for proteomic analysis.

### Mass spectrometry data analysis and label-free quantitation

Sample preparation and mass spectrometry analysis was performed similar to previously described work^13-16^. Precipitated proteins were re-dissolved in 1% sodium 3-[(2-methyl-2-undecyl-1,3-dioxolan-4-yl)methoxy]-1-propanesulfonate (commercially available as RapiGest, Waters), 50 mM Tris-HCl pH 8.0, 1 mM TCEP. Cysteines were alkylated by addition of 20 mM Iodoacetamide and incubation for 20 min at 25°C in the dark and the reaction quenched by addition of 20 mM DTT. Samples were diluted to 0.1% Rapigest with 50 mM Tris-HCl pH 8.0 and Trypsin (sequencing grade, Promega) was added at a 1:50 ratio. Proteins were digested overnight at 37°C under constant shaking.

Samples from four biological replicates (0.5 μg of digest for the secretome analyses) were injected in an interleaved manner onto a 2 cm x 100 μm trap column and separated on a 50 cm x 75 μm EasySpray Pepmap C18 reversed-phase column (Thermo Fisher Scientific) on a Dionex 3000 Ultimate RSLC. Peptides were eluted by a linear 3-hour gradient of 95% A/5% B to 35% B (A: H_2_O, 0.1% Formic acid (FA); B: 80% ACN, 0.08% FA) at 300 nl/min into a LTQ Orbitrap Velos (Thermo-Fisher Scientific). Data was acquired using a data-dependent “top 20” method, dynamically choosing the most abundant precursor ions from the survey scan (400-1600 Th, 60,000 resolution, AGC target value 10^6^). Precursors above the threshold of 2000 counts were isolated within a 2 Th window and fragmented by CID in the LTQ Velos using normalized collision energy of 35 and an activation time of 10 ms. Dynamic exclusion was defined by a list size of 500 features and exclusion duration of 60 s. Lock mass was used and set to 445.120025 for ions of polydimethylcyclosiloxane (PCM).

Label-free quantitation was performed using MaxQuant 1.5.7.4^17^. Data were searched against the Uniprot database of *S. aureus* NCTC8325 (downloaded on 29.03.17) containing 2,889 sequences and a list of common contaminants in proteomics experiments using the following settings: enzyme Trypsin/P, allowing for 2 missed cleavage, fixed modifications were carbamidomethyl (C), variable modifications were set to Acetyl (Protein N-term), Deamidation (NQ) and Oxidation (M). MS/MS tolerance was set to 0.5 Da, precursor tolerance was set to 6 ppm. Peptide and Protein FDR was set to 0.01, minimal peptide length was 7, and one unique peptide was required. Re-quantify and retention time alignment (2 min) were enabled. If no intensities were detected in one condition and the other condition had intensities in at least in 3 out of 4 replicates, values were imputed in Perseus v1.5.1.1 using default parameters^18^. A student’s t-test (two-tailed, homoscedastic) was performed on the LFQ intensities and only proteins with p<0.05 and a fold-change >2-fold were considered significant.

### Data availability statement

Raw mass spectrometry data that support the findings of this study have been deposited to the ProteomeXchange Consortium via the PRIDE^19^ partner repository with the dataset identifier PXD011358. All other data supporting the findings of this study are available within the paper and its supplementary information files.

### Bacterial membrane potential detection

To assess bacterial membrane potential, the method of Miyata *et al*.^20^ was adapted, using the BacLight bacterial membrane potential kit (Invitrogen). In brief, overnight cultures of *E. coli* MG1655 harboring pBAD18-Cm derivatives were sub-cultured into LB medium containing appropriate antibiotics to an OD_600_ of 0.1 and cultured aerobically to OD_600_ of 0.5, before supplementation with 0.2% L-arabinose. Cells were cultured for a further hour then diluted to 1 × 10^6^ cells per ml in sterile PBS, and a 1ml aliquot of bacterial suspension was added to each flow cytometry tube. For the depolarized control, 25 μL of 500 μM carbonyl cyanide m-chlorophenyl hydrazone (CCCP), provided in the BacLight bacterial membrane potential kit, was added. Next, 3 μL of 3mM DiOC_2_(3) was added to each sample, which was mixed and incubated at room temperature for 30 minutes. Cells were subsequently sorted using an LSRFortessa cell analyzer (BD Biosciences, San Jose, CA). The DiOC_2_(3) dye was excited at 488 nm, with fluorescent emissions detected using Alexa488 (Ex 488nm, Em 530/30 nm) and Alexa568 (Ex 561 nm, Em 610/20 nm). In total, 20000 events were collected for each sample, with forward- and side-scatter parameters used to gate the bacteria. The forward scatter, side scatter, and fluorescence were collected with logarithmic signal amplification. The gated populations were then analyzed with FlowJo software by generating a dot plot of red versus green fluorescence.

To assess changes in membrane potential and permeabilization, the same *E. coli* strains were grown as described above. An ‘uninduced’ sample of *E. coli* harboring each of pBAD18-Cm (empty), pAmiAss-TspA_CT_ and pAmiAss-TspA_CT_-TsaI was collected and adjusted to OD_600_ of 0.2 before the addition of 2 µM DiSC_3_(5) and 200 nM Sytox Green. A permeabilized control sample of cells harboring empty vector and containing 10 μg/ml Polymyxin B was also prepared and supplemented with both dyes. All samples were then incubated at 37°C for 5 minutes before being analyzed by microscopy. To induce protein production from the pBAD18-Cm vector, cell suspensions were adjusted to OD_600_ of 0.2 and supplemented with 0.2% L-arabinose for the indicated period of time (10-60 min) before incubating with DiSC_3_(5) and Sytox Green, as above. Imaging of DiSC_3_(5) and Sytox Green stained cells was carried out Nikon Eclipse Ti equipped with Sutter Instrument Lambda LS light source, Nikon Plan Apo 100×/1.40 NA Oil Ph3 objective, and Photometrics Prime sCMOS, and Cy5 and GFP filters, respectively. The images were captured using Metamorph 7.7 (Molecular Devices) and analyzed using ImageJ. For the analysis, the phase contrast images acquired in parallel to the fluorescence images were used to identify cells as regions of interest, for which average DiSC_3_(5) and Sytox Green fluorescence intensity was measured from the corresponding background-subtracted fluorescence images. Data was then plotted as a scatter plot for DiSC_3_(5) and Sytox Green fluorescence with each point representing an individual cell.

### Mouse pneumonia model

*S. aureus* RN6390 (WT) or the isogenic Δ*essC* strain were subcultured at 1:100 dilution from an overnight culture into fresh TSB medium. Cells were grown at 37°C with shaking until an OD_600_ of 0.5 was reached, before harvesting and washing three times in 1x PBS. Cells were finally resuspended in 1 x PBS to 1.2 × 10^10^ cfu/ml. Female 10-12 week old C57/B6 J mice were purchased from Charles River U.K.. Mice were acclimatized for a 10 day period prior to starting the experiment. Mice were randomized to cages and treatment groups, and analyses were performed blind. Mice were anaesthetized (gaseous isofluorane) and infected intranasally with 25 μL of the bacterial suspension to give a final infected dose of 3 × 10^8^ cfu per mouse. At 24 hours post infection the lungs and livers were harvested and the bacterial load determined by plating serial dilutions of tissue homogenates.

### Quantitative reverse transcription polymerase chain reaction (qRT-PCR)

For RNA was harvested from 10 snap-frozen zebrafish larvae infected with *S. aureus* at 6 hpi using the RNeasy Mini Kit (Qiagen) as per manufacturer’s instructions. RNA was reverse transcribed into cDNA using QuantiTect Reverse Transcription Kit (Qiagen) as per manufacturer’s instructions with 500 ng of RNA. Quantitative RT-PCR reactions were performed on four biological replicates, each with two technical replicates. For qRT-PCR reactions, 50 ng of cDNA was used per reaction with SYBR Green Reaction Mix (Thermo Fisher Scientific) on a Rotor GeneQ (Qiagen) thermocycler. Oligonucleotides for *il-1b, cxcl8* and *eef1a1a* are listed in *SI Appendix*, Table S4. Quantities of cDNA were normalised using the housekeeping gene *eef1a1a* and the 2-ΔΔCT method was used for analysis^21^.

### Imaging of *S. aureus* – leukocyte interactions *in vivo*

To observe *S. aureus* – leukocyte interactions, Tg(*lyz*::dsRed)^nz50^ and Tg(*mpeg1*::Gal4-FF)^gl25^/Tg(*UAS*-*E1b*::*nfsB*.mCherry)^c264^ zebrafish larvae were infected with *S. aureus* strains chromosomally labelled with GFP. To follow neutrophil and macrophage recruitment, infected larvae were anaesthetised with 200 μg/ml Tricaine and the HBV imaged at 0, 3, and 6 hpi by fluorescence stereomicroscopy and multiple position z stacks of up to 400 μm were acquired using a Leica M205FA stereomicroscope with a 10x (NA 0.5) dry objective. Neutrophil and macrophage quantifications were performed manually throughout the individual z stacks using Fiji – ImageJ (ver 1.0).

### Statistical analysis

For statistical analysis GraphPad Prism 6.0 software was used. In survival assays statistical analysis was done using a Log rank (Mantel-Cox) test. To analyse bacterial kinetics, CFU counts were Log10 transformed and the significance between two independent groups was determined by an unpaired *t* test. When more than two groups were compared, significance was determined by using one-way ANOVA with Sidak’s comparison. Gene expression levels were quantified on Log2 data and significance was determined by using one-way ANOVA with Sidak’s Multiple Comparison test. For leukocyte cell counts analysis (non-parametric data), significance between multiple selected groups was determined using Kruskal-Wallis test with Dunn’s correction. Significance is indicated ns, non-significant, *, p ≤0.05; **, p ≤ 0.01; ***, p ≤ 0.001; ****, p ≤ 0.0001.

---

**Figure S1.**
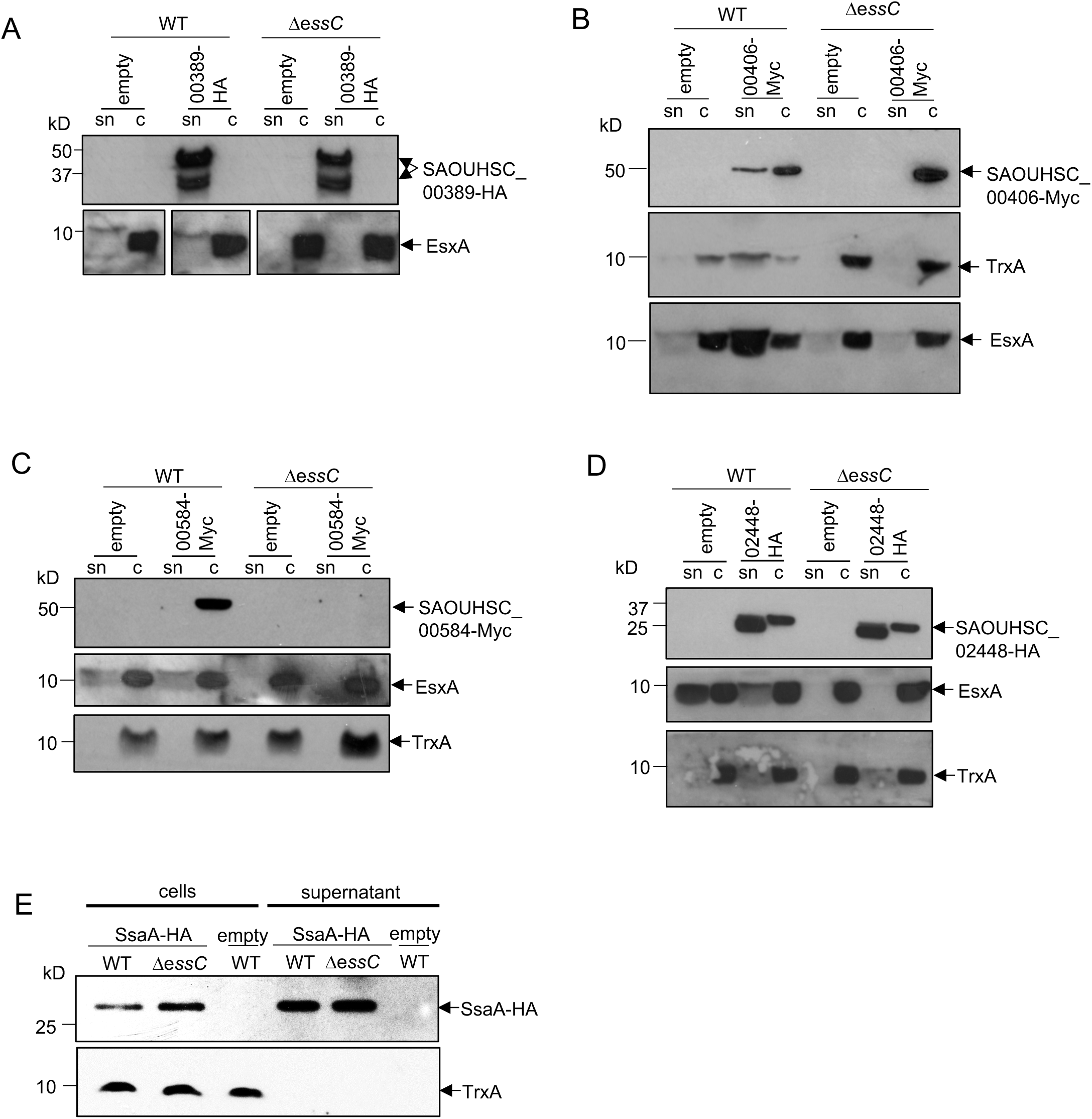
HA-tagged variants of SAOUHSC_00389, SAOUHSC_02448 and SsaA are not secreted by the T7SS. The wild type RN6390 and Δ*essC* mutant carrying A-D pRAB11 (empty) or pRAB11 encoding A. C-terminally HA-tagged SAOUHSC_00389; B. C-terminally Myc-tagged SAOUHSC_00406; C. C-terminally Myc-tagged tagged SAOUHSC_00584; or D. C-terminally HA-tagged SAOUHSC_02448, or E. pRMC2 (empty) or encoding C-terminally HA-tagged SsaA were cultured in TSB medium and supplemented with 200ng/ml ATc at OD_600_ of 0.5. When cultures reached OD_600_ of 2.0, samples were withdrawn and separated into culture supernatant (sn) and cellular (c) fractions. Samples were separated on 12% bis-Tris gels and immunoblotted with anti-HA, anti-Myc, anti-EsxA or anti-TrxA antibodies, as indicated.

**Figure S2.**
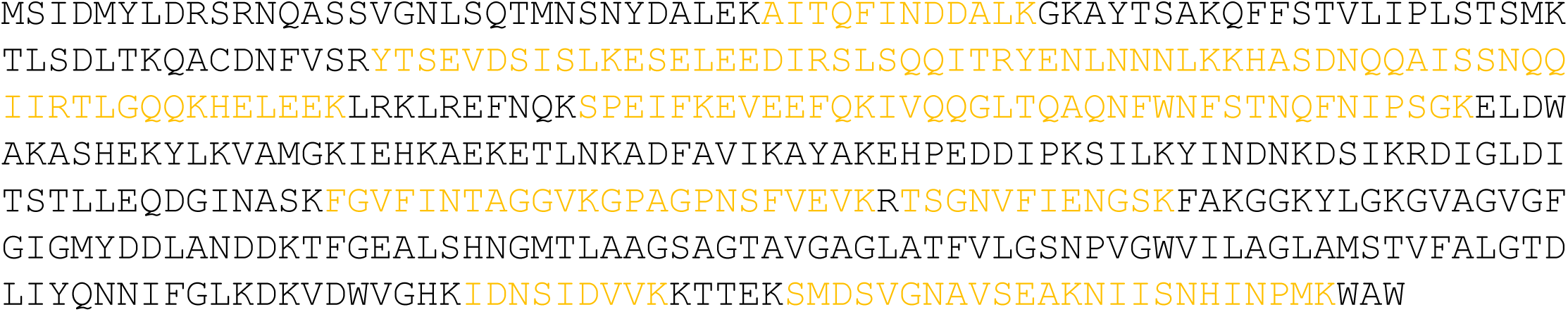
Peptide coverage of TspA from proteomic analysis. Regions of TspA coloured orange were detected by mass spectroscopy analysis.

**Figure S3.**
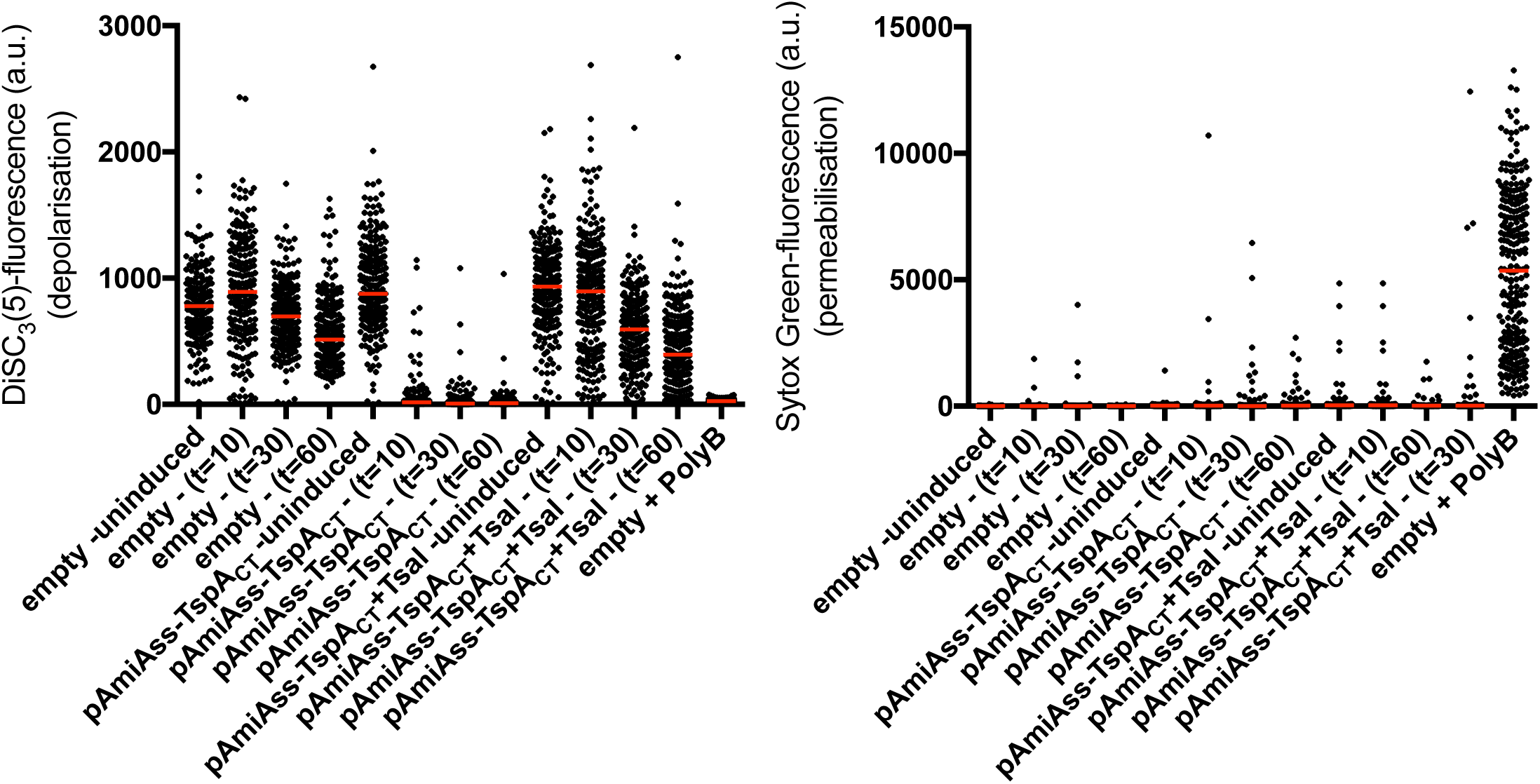
Rapid membrane depolarization by the C-terminal domain of TspA. Fluorescence intensity changes in DiSC_3_(5) (left) and Sytox Green (right) in uninduced cells harbouring pBAD18-Cm (empty), pBAD18-AmiAss-TspA_CT_ and pBAD18-AmiAss-TspA_CT_ and upon induction with 0.2% L-arabinose. Cells were then induced for 10, 30 and 60 minutes before samples removed and cells incubated with DiSC_3_(5) and Sytox Green. Fluorescence intensity of DiSC_3_(5) and Sytox Green also measured upon addition of Polymixin B to empty vector as a control for membrane depolarization and permeabilisation.

**Figure S4.**
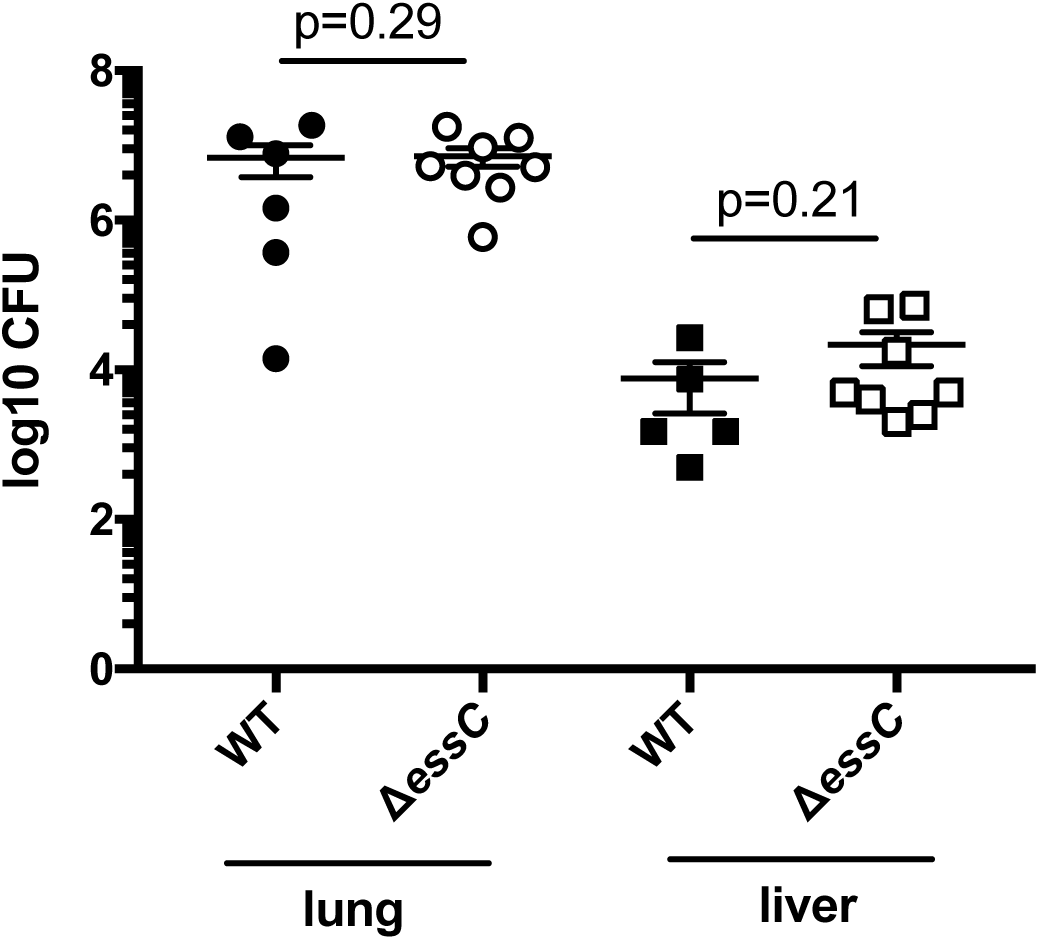
No difference in bacterial burden between wild type and *essC* mutant strains in a 24 hour murine pneumonia infection model. Female 10-12 week old C57/B6 mice were challenged with 3 × 10^8^ cfu/ml of RN6390 (WT) or the isogenic Δ*essC* strain. Bacterial load was determined in liver and lungs 24 hours after infection. Mean ± SEM (horizontal bars) is shown. Significance testing performed by unpaired *t* test.

**Figure S5.**
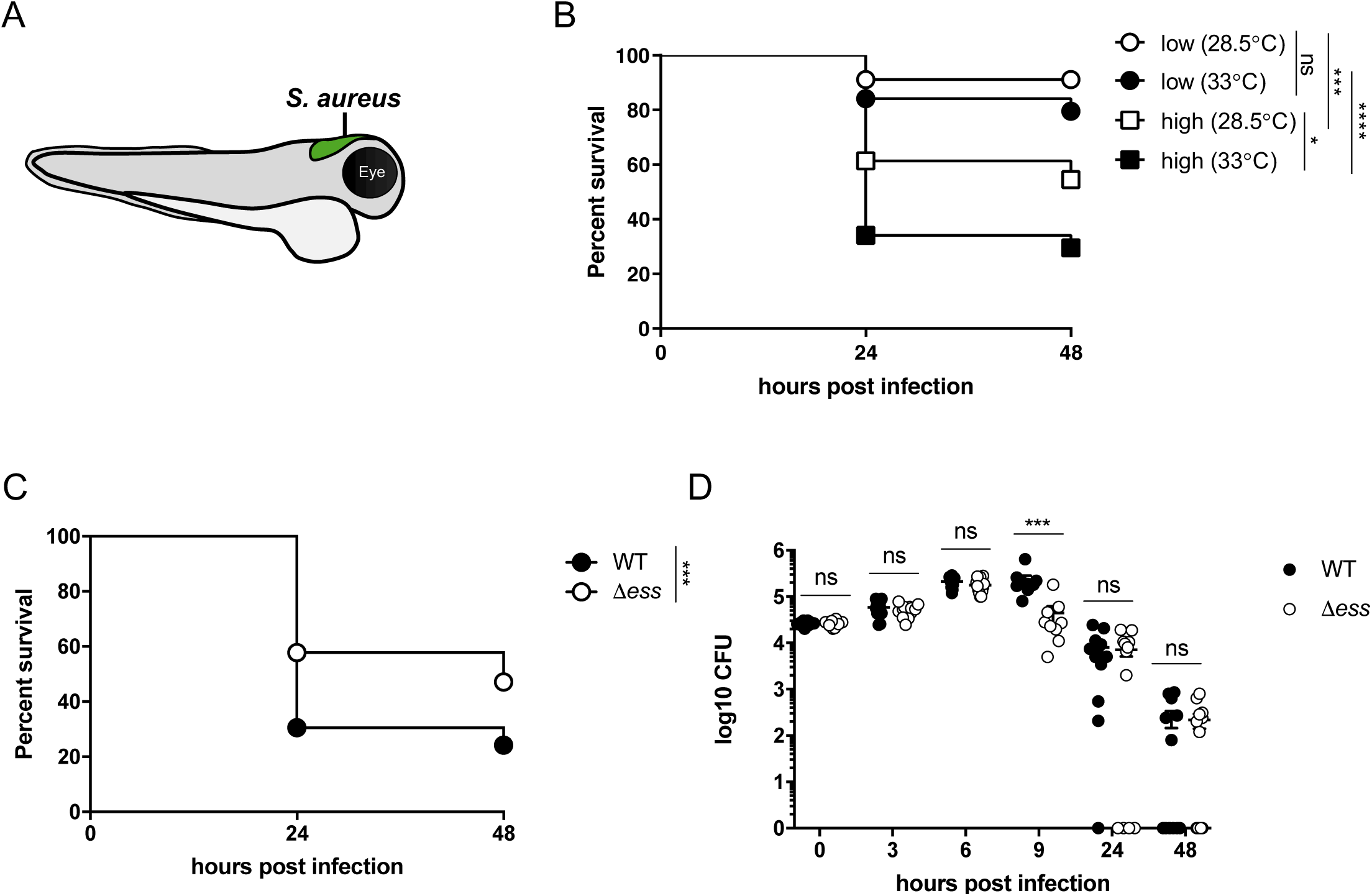
Developing a zebrafish model to assess T7SS activity *in vivo*. A. Schematic of zebrafish larvae showing the site of *S. aureus* injection into the hindbrain ventricle. B. Survival curves of *lyz*:dsRed zebrafish larvae injected with wild type RN6390 chromosomally tagged with GFP. Zebrafish were injected at 3 dpf with a low (∼7 × 10^3^ cfu), medium (∼1.4 × 10^4^ cfu) or high (∼2 × 10^4^ cfu) dose of RN6390-gfp, incubated at 28.5°C or 33°C and monitored for 48 hpi. Data are pooled from two independent experiments (*n*=22-25 larvae per experiment). Results are plotted as a Kaplan-Meier survival curve and the p value between conditions was determined by log-rank Mantel-Cox test. C. Survival curves of *lyz*:dsRed larvae infected in the hindbrain 3 dpf with RN6390-gfp or RN6390 Δ*ess*-gfp at a dose of ∼2 × 10^4^ cfu and incubated at 33°C for 48 hpi. Data are pooled from three independent experiments (*n*=28-50 larvae per experiment. Results are plotted as a Kaplan-Meier survival curve and the p value between conditions was determined by log-rank Mantel-Cox test. D. Enumeration of recovered bacteria at 0, 3, 6, 9, 24 or 48 hpi from zebrafish larvae infected with RN6390-gfp or RN6390 Δ*ess*-gfp. Pooled data from 3 independent experiments. Circles represent individual larva, and only larvae having survived the infection were included. Mean ± SEM also shown (horizontal bars). Significance was tested using an unpaired *t* test. \**p*<0.05 \*\**p*<0.01, *** *p*<0.001, **** *p*<0.0001, ns, not significant.

**Figure S6.**
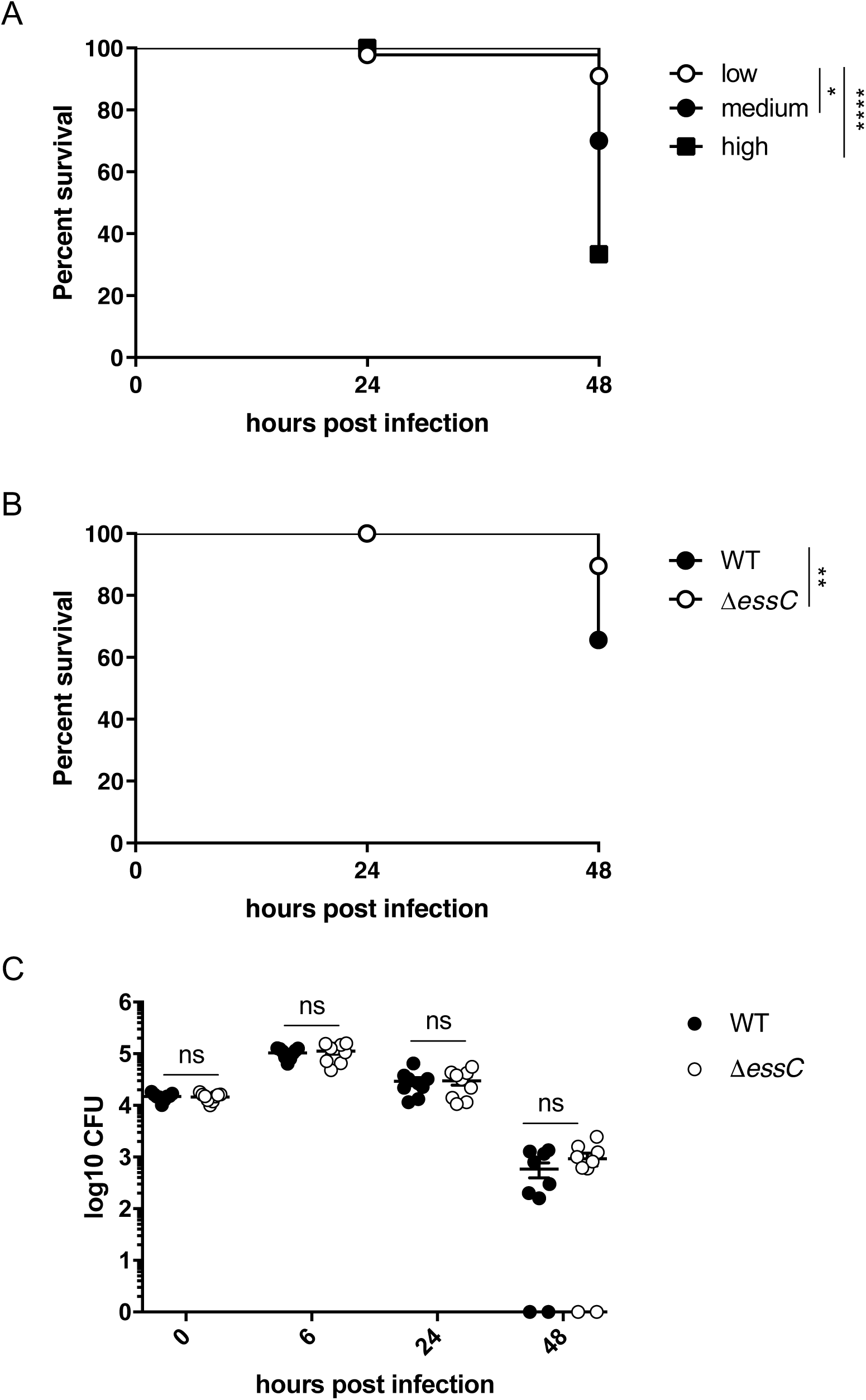
*S. aureus* COL also exhibits dose and T7SS-dependent zebrafish mortality. A. Survival curves of WT zebrafish larvae injected with wild type COL chromosomally tagged with mCherry. Zebrafish were injected at 3 dpf with a low (∼7 × 10^3^ cfu), medium (∼1.5 × 10^4^ cfu) or high dose (∼2 × 10^4^ cfu) of COL-mCherry, incubated at 33°C and monitored for 48 hpi. Data are pooled from three independent experiments. Results are plotted as a Kaplan-Meier survival curve and the *p* value between conditions was determined by log-rank Mantel-Cox test. B. Survival curves of zebrafish larvae injected in the hindbrain 3 dpf with COL-mCherry (WT) or COL Δ*essC*-mCherry at a dose of ∼1.6 × 10^4^ cfu and incubated at 33°C for 48 hpi. Data are pooled from three independent experiments (*n*=23-30 larvae per experiment). Results are plotted as Kaplan-Meier survival curves and the *p* value between conditions was determined by the log-rank Mantel Cox test. C. Enumeration of recovered bacteria at 0, 6, 24 or 48 hpi from zebrafish larvae infected with COL-mCherry (WT) or COL Δ*essC*-mCherry. Circles represent individual larvae and data pooled data from 3 independent experiments. Mean ± SEM also shown (horizontal bars). Significance was tested using an unpaired *t* test. \**p*<0.05 \*\**p*<0.01, *** *p*<0.001, **** p<0.0001, ns, not significant.

**Figure S7.**
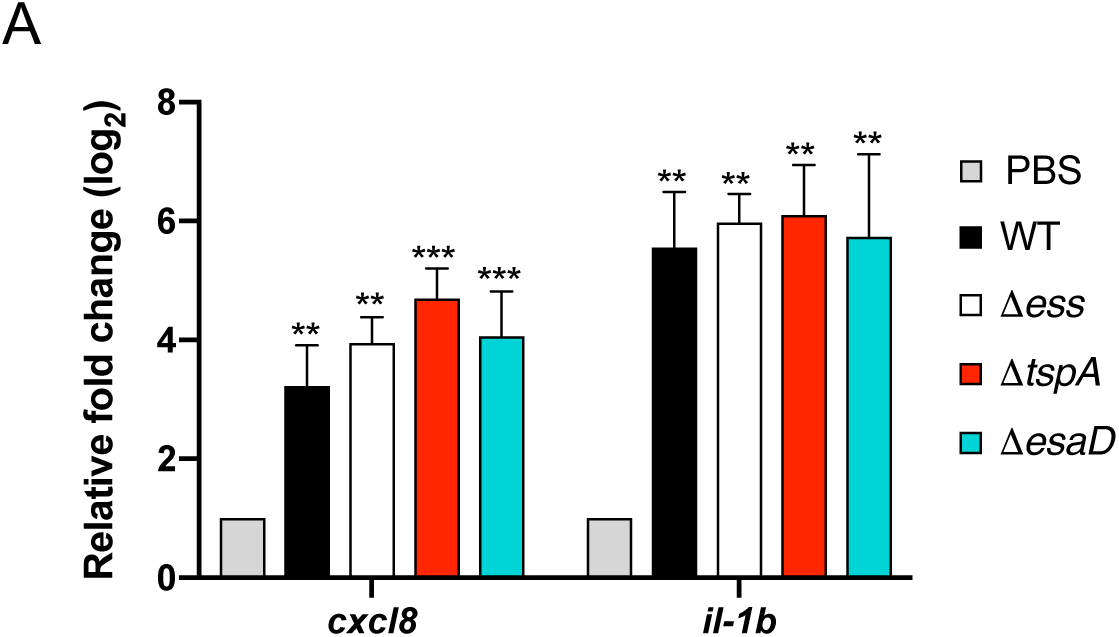
*S. aureus* infection elicits a strong inflammatory response independent of the T7SS. A. *S. aureus* RN6390-gfp, RN6390 Δ*ess*-gfp, Δ*tspA*-gfp or RN6390 Δ*esaD*-gfp were injected in 3 dpf zebrafish larvae at a dose of ∼2 × 10^4^ CFU and incubated at 33°C. Expression of *cxcl8* and *il-1b* was determined at 6 hpi when the bacterial burden between among strains was similar. Mean relative *cxcl8* and *il-1b* gene expression levels (qRT-PCR) were quantified and values were normalised to the PBS-injected larvae. Therefore, the *p* value (indicated above the bars) represents the statistical significance of the indicated *S. aureus* strains in comparison to the PBS-injected larvae. Pooled data from 4 independent experiments where 10 larvae were sacrificed after 6 hpi. Error bars represent mean with SEM (horizontal bars). Significance was performed using a one-way ANOVA with Sidak’s correction. \**p*<0.05 \*\**p*<0.01, *** *p*<0.001, **** *p*<0.0001, ns, not significant.

**Figure S8.**
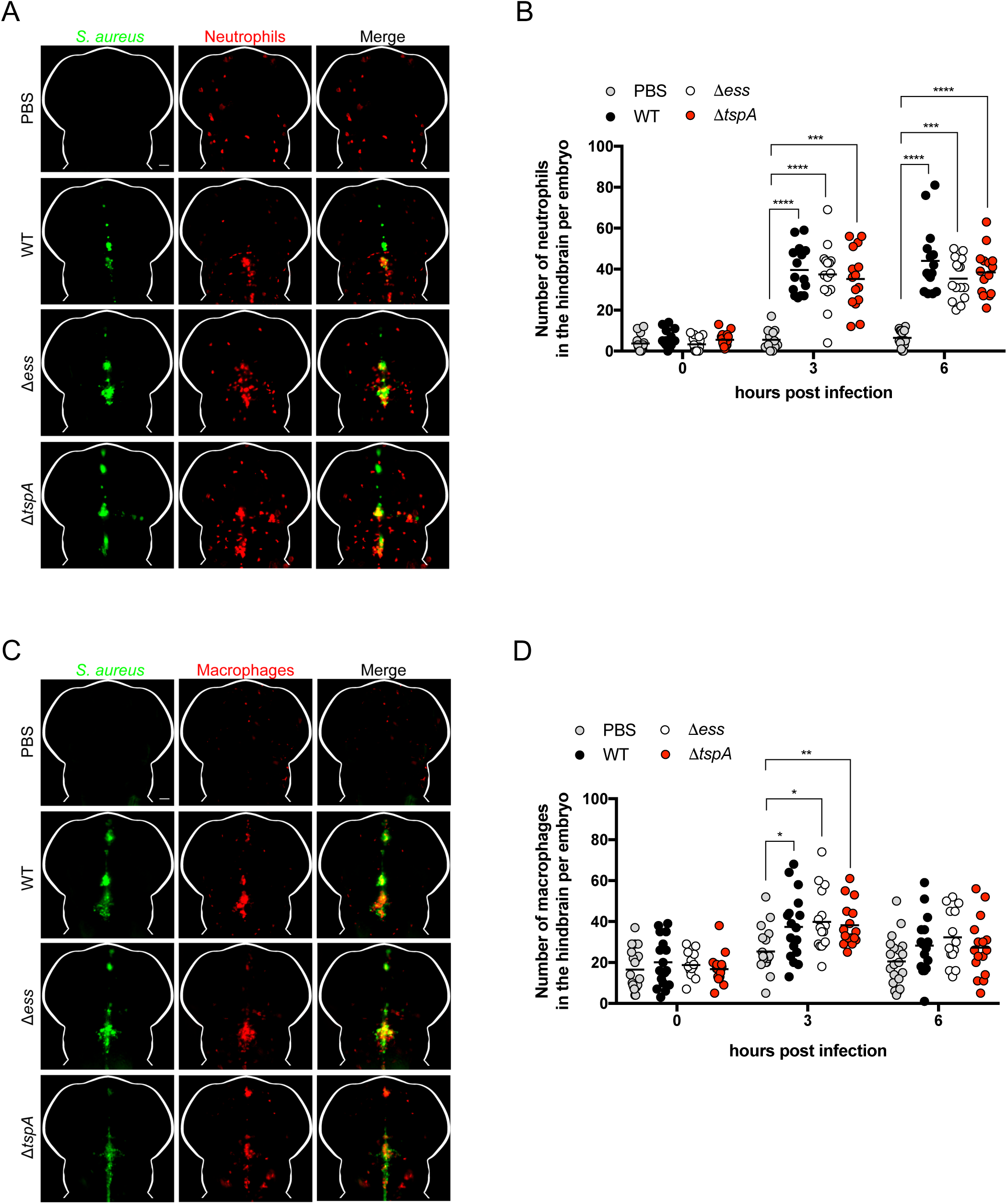
T7SS independent leukocyte recruitment to *S. aureus* infection. A+B. Neutrophils were imaged and counted in the whole hindbrain for Tg(*lyz*::dsRed) larvae and C+D. Macrophages were imaged and counted in the whole hindbrain for Tg(*mpeg1*::G/U::mCherry) larvae injected with PBS, RN6390-gfp, RN6390 Δ*ess*-gfp or RN6390 Δ*tspA*-gfp. Larvae were imaged using a fluorescent stereomicroscope at 0, 3 and 6 hpi and data obtained from three independent experiments with 5-6 larvae imaged per strain. Data points represent an individual larvae with the geometric mean. Significance testing performed using Kruskal-Wallis test with Dunn’s correction. \**p*<0.05 \*\**p*<0.01, *** *p*<0.001, **** *p*<0.0001, ns, not significant. Representative images of a single z stack of neutrophil or macrophage recruitment at 6 hpi is shown in A+C, respectively. Leukocytes are labelled in red and S. aureus in green, with overlay in yellow. Scale bar = 50 μm Scale bar Scale bar = 50 μm.

**Figure S9.**
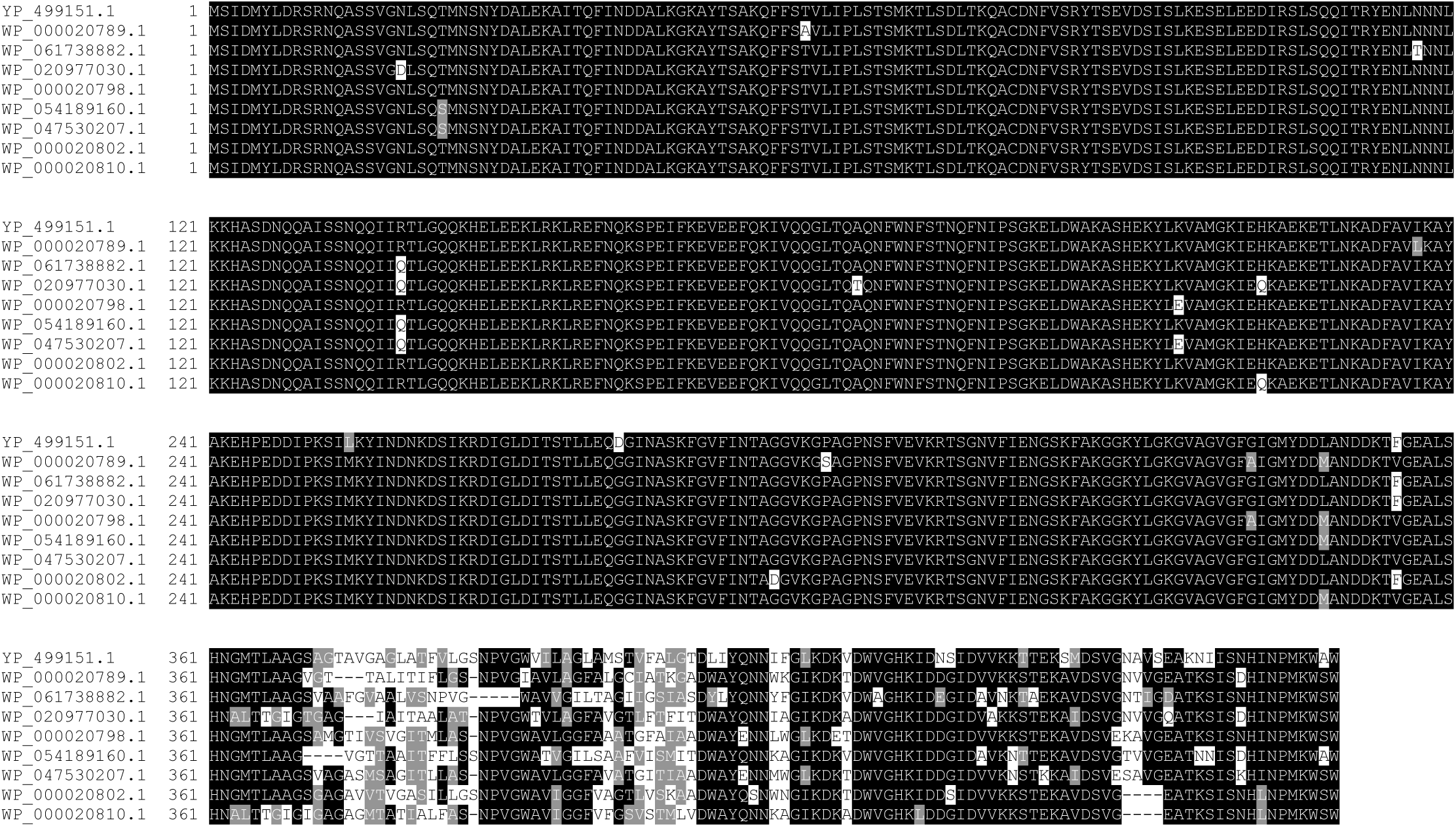
TspA proteins encoded by *S. aureus* strains show variability within the channel-forming domain. A selection of TspA protein sequences encoded by *S. aureus* strains were extracted from NCBI (ncbi.nlm.nih.gov/), aligned using ClustalW (http://www.ch.embnet.org/software/ClustalW.html) and shaded with Boxshade (http://www.ch.embnet.org/software/BOX_form.html).

