## Supplemental Tables 2,3 and 4 for "A membrane-depolarizing toxin substrate of the *Staphylococcus aureus* Type VII secretion system mediates intra-species competition"

**Table S2**. Strains used in this study

| Strain | Relevant genotype or description | Source or reference |
|---|---|---|
| *S. aureus* strains | | |
| RN6390 | NCTC8325 derivative, *rbsU*, *tcaR*, cured of φ11, φ12, φ13 | Reference[1] |
| RNΔ*essC* | As RN6390, Δ*essC* | Reference[2] |
| RNΔ*ess* | Complete deletion from *esxA* – *esaG* | Reference[2] |
| RNΔ*esaD* | As RN6390, Δ*esaD* | Reference[3] |
| RNΔ*tspA* | As RN6390, Δ*tspA* (*saouhsc00584)* | This work |
| RNΔ*saouhsc00268-00278* | As RN6390, Δ*esaD-saouhsc00278* | Reference[3] |
| RNΔ*saouhsc00585-00602* | As RN6390, Δ*saouhsc00585-saouhsc00602* | This work |
| FRU1 | RN6390 Δ*saouhsc00268-00278*, Δ*saouhsc00585-00602* | This work |
| RN6390*::ermC* | As RN6390, with *ermC* resistance gene chromosomal insertion | Reference[3] |
| RNΔ*saouhsc00268-00278::ermC* | As RN6390, Δ*esaD-saouhsc00278* with *ermC* resistance gene from RN6390*::ermC* (phage ϕ11 transduction) | This work |
| FRU1*::ermC* | RN6390 Δ*saouhsc00268-00278*, Δ*saouhsc00585-00602* with *ermC* resistance gene from RN6390*::ermC* (phage ϕ11 transduction) | This work |
| RN6390::GFP | As RN6390, with markerless GFP insertion | This work |
| RNΔ*saouhsc00268-00278::gfp* | As RNΔ*esaD-saouhsc00278,* with markerless GFP insertion | RNΔ*saouhsc00268-00278* |
| RNΔ*saouhsc00585-00602::gfp* | As RNΔ*saouhsc00585-00602*, with markerless GFP insertion | This work |
| FRU1*::gfp* | RN6390 Δ*saouhsc00268-00278*, Δ*saouhsc00585-00602*, with markerless GFP insertion | This work |
| COL | MRSA, *agr* | Reference[4] |
| COLΔ*essC* | As COL, Δ*essC* | This work |

| | | |
|---|---|---|
| COLΔ*esaD* | As COL, Δ*esaD* | This work |
| COLΔ*tspA* | As COL, Δ*tspA* (*sacol0643*) | This work |
| COL*::mCherry* | As COL, with markerless mCherry insertion | This work |
| COLΔ*essC::mCherry* | As COL, Δ*essC* with markerless mCherry insertion | This work |
| COLΔ*esaD::mCherry* | As COL, Δ*esaD* with markerless mCherry insertion | This work |
| COLΔ*tspA::mCherry* | As COL, Δ*tspA* (*sacol0643*) with markerless mCherry insertion | This work |
| *E. coli* strains | | |
| JM110 | *rpsL thr leu thi lacY galK galT ara tonA tsx dam dcm glnV44 Δ(lac-proAB)* e14- [F' *traD36* $proAB^{+}$ $lacI^{q}$ *lacZ*ΔM15] *hsdR17*($rK^{-}mK^{+}$) | Stratagene |
| MG1655 | *E. coli K-12, F–, λ–, ilvG, rfb-50, rph-1* | Reference[5] |
| SG3000 | As MG1655, Δ*tatABCD* | Reference[6] |

**Table S3**. Plasmids used in this study

| Plasmid | Relevant genotype or description | Source or reference |
|---|---|---|
| pIMAY | *E. coli/S. aureus* shuttle vector, temperature sensitive, cml$^r$ | Reference[7] |
| pIMAY-esaD | pIMAY carrying *esaD* deletion allele | Reference[3] |
| pIMAY-essC | pIMAY carrying *essC* deletion allele | Reference[2] |
| pIMAY-saouhsc00268-00278 | pIMAY carrying *esaD-saouhsc00278* deletion allele | Reference[3] |
| pIMAY-tspA | pIMAY carrying *tspA* (*saouhsc00584*) deletion allele | This work |
| pIMAY-saouhsc00585-00602 | pIMAY carrying *saouhsc00585-saouhsc00602* deletion allele | This work |
| pTH100 | Plasmid for markerless integration of GFP into *S. aureus* | Reference[8] |
| pRN111 | Plasmid for markerless integration of mCherry into *S. aureus* | Reference[8] |
| pBAD18-cm | Glucose-repressible/arabinose inducible vector; cml$^r$ | Reference[9] |
| pBAD18-TspA | pBAD18-Cm producing TspA | This work |
| pBAD18-TspA$_{CT}$ | pBAD18-Cm producing amino acids 218-469 of TspA | This work |
| pBAD18-AmiAss-TspA$_{CT}$ | pBAD18-Cm producing *E. coli* AmiA signal sequence fused to amino acids 218-469 of TspA. | This work |
| pBAD18-AmiAss-TspA$_{CT}$ + TsaI | As pBAD18-AmiAss-TspA$_{CT}$ but also producing TsaI | This work |
| pSU-PROM | Cloning vector for expression of genes under the control of the *tat* promoter; Km$^R$ | Reference[10] |
| pSU-PROM-TsaI | pSU-PROM producing TsaI | This work |
| pRAB11 | *E. coli/S. aureus* shuttle vector, inducible protein expression, amp$^r$, cml$^r$ | Reference[11] |
| pRAB11-TspA-Myc | pRAB11 producing C-terminally Myc-tagged TspA | This work |
| pRAB11-02448-HA | pRAB11 producing C-terminally HA-tagged SAOUHSC_02448 | This work |
| pRAB11-00406-Myc | pRAB11 producing C-terminally Myc-tagged SAOUHSC_00406 | This work |
| pRAB11-00389-HA | pRAB11 producing C-terminally HA-tagged SAOUHSC_00389 | This work |
| pRAB11-00585-HA | pRAB11 producing C-terminal HA-tagged SAOUHSC_00585 | This work |
| pRMC2 | *E. coli-S. aureus* shuttle vector, inducible protein expression, amp$^r$, cml$^r$ | Reference[12] |
| pRMC2 -SsaA-HA | pRMC2 producing C-terminally HA-tagged SsaA | This work |

**Table S4.** Oligonucleotides and cloning strategies used in this study

| Primer | Nucleotide Sequence (5'-3') |
|---|---|
| TspA A1 | TAGGTACCGCTAATACATGCACGGC |
| TspA A2 | CGCCCATTTCATCGGCATGCTCCTTTTC |
| TspA B1 | ATGCCGATGAAATGGGCGTGGTGAGTT |
| TspA B2 | CCGGTACCTGCTTTTTAAGTTTGGCATA |
| TspA out1 | TCGCAAAGCAATATCCAC |
| TspA out2 | TTGGTACCAATGGGGCATTTACGA |
| tspA cmyc fw | GCGCGGTACCAGGAGGTTTCTAGTTATGAGTATTGACATGTATTTAG AC |
| tspA cmyc rv | GCGCGAGCTCTCACAGATCCTCTTCTGAGATGAGTTTTTGTTCCCAC GCCCATTTCATTGGATTTATATG |
| 02448 cha fw | GCGCGGTACCAGGAGGTTTCTAGTTATGGGAGTTAAAAGTGTG |
| 02448 cha rv | GCGCGAGCTCTTATGCATAATCTGGAACATCATATGGATATTTTTTC C ATAAGAAGTC |
| 00406 cmyc fw | GCGCGGTACCAGGAGGTTTCTAGTTATGTTGAGTAGGAAG |
| 00406 cmyc rv | GCGCGAGCTCTTACAGATCCTCTTCTGAGATGAGTTTTTGTTCTAGT A ATCCACCTATTTGTG |
| 00389 cha fw | GCGCGGTACCAGGAGGTTTCTAGTTATGAAAATAACAACGATTGC |
| 00389 cha rv | GCGCGAGCTCTTATGCATAATCTGGAACATCATATGGATATTTTATAT TCACTTCAATG |
| 00585 cha fw | GCGC AGATCT AGGAGG TTT CTA GTT ATG TTT TTA ATA TTA AGG TT |
| 00585 cha rv | GCGCGAATTCTTAAGCATAATCTGGAACATCATATGGATATTGCTTTT TAAGTTTGGCATAAAC |
| pRMC2-ssaA-bglII-for | GGAGATCTAGAGTGTTTTGATTATTGGGA |
| pRMC2- SSaA-HA-rev-EcoRI | GGGAATTCTTATGCATAATCTGGAACATCATATGGATAATGAATGAA ATTATATGAACC |
| AmiAss fw | GCGCGCTAGCCAGAGGAGGAGCCATGAGCACTTTTAAACCAC |
| AmiAss rv | GCGCGGTACCGTCTTTGGCGATGGCTTGCGAC |
| tspA fl fw | GCGCTCTAGACAGAGGAGGAGCCATGAGTATTGACATGTATTTAGA C |
| tspA fl rv | GCGCGTCGACTTACCACGCCCATTTCATTGG |
| tspA cp fw | GCGCTCTAGACAGAGGAGGAGCCATGATTGAACATAAAGCAGAGAA AG |
| tspA pp fw | GCGCTCTAGAATTGAACATAAAGCAGAGAAAG |
| tsaI fw | GCGCGTCGACCAGAGGAGGAGCCATGCTTTTTAATATTAAGG |
| tsaI rv | GCGCGCATGCTTATTGCTTTTTAAGTTTGGCATAAAC |
| pSU-tsaI fw | GCGC GGATCC ATG CTT TTT AAT ATT AAG G |
| pSU-tsaI rv | GCGC CTCGAG TTATTG CTT TTT AAG TTT GGC |
| saouhsc00585-00602 A1 | GCGCGAATTCCAGGTGGAGTGAAAGGCCCAGC |
| saouhsc00585-00602 A2 | CTATTGGTTTTTATTAAAAAGCAAAACTCACCAC |

| | |
|---|---|
| saouhsc00585-00602 B1 | TTGCTTTTTAATAAAAACCAATAGAAATTACCAA |
| saouhsc00585-00602 B2 | GCGCGAGCTCCCGGTTGATTGTTCTGATGTAC |
| saouhsc00585-00602 out1 | GCATATGCCAAAGAACATCCAG |
| saouhsc00585-00602 out2 | GCTCTTGTAATGCTGCAACTGC |
| *eef1a1a* for | AAGCTTGAAGACAACCCCAAGAGC |
| *eef1a1a* rev | ACTCCTTTAATCACTCCCACCGCA |
| *cxcl8* for | TGTGTTATTGTTTTCCTGGCATTTC |
| *cxcl8* rev | GCGACAGCGTGGATCTACAG |
| *il1b* for | GAACAGAATGAAGCACATCAAACC |
| *il1b* rev | ACGGCACTGAATCCACCAC |
